# Open syntaxin overcomes synaptic transmission defects in diverse *C. elegans* exocytosis mutants

**DOI:** 10.1101/2020.01.10.901835

**Authors:** Chi-Wei Tien, Bin Yu, Mengjia Huang, Karolina P. Stepien, Kyoko Sugita, Xiaoyu Xie, Liping Han, Philippe P. Monnier, Mei Zhen, Josep Rizo, Shangbang Gao, Shuzo Sugita

**Affiliations:** Division of Fundamental Neurobiology, Krembil Brain Institute, University Health Network, Ontario, M5T 2S8, Canada; Department of Physiology, Faculty of Medicine, University of Toronto, Toronto, Ontario, M5S 1A8, Canada; College of Life Science and Technology, Huazhong University of Science and Technology, Wuhan, China, 430074; Department of Biophysics, University of Texas Southwestern Medical Center, Dallas, United States; Department of Biochemistry, University of Texas Southwestern Medical Center, Dallas, United States; Department of Pharmacology, University of Texas Southwestern Medical Center, Dallas, United States; Division of Genetics and Development, Krembil Brain Institute, University Health Network, Ontario, M5T 2S8, Canada; Department of Anesthesiology, Dalian Medical University, Dalian, Liaoning, China; Department of Anesthesiology, Dalian Municipal Friendship Hospital, Dalian Medical University, Dalian, Liaoning, China; Department of Ophthalmology & Vision Sciences, University of Toronto, Toronto, Ontario, M5S 1A8, Canada; Lunenfeld-Tanenbaum Research Institute, Mount Sinai Hospital, Toronto, Ontario, M5G 1X5, Canada; Department of Molecular Genetics, Faculty of Medicine, University of Toronto, Toronto, Ontario, M5S 1A8, Canada

**Keywords:** exocytosis, vesicle priming, SNARE, synaptotagmin, Munc18/UNC-18, Munc13/UNC-13

## Abstract

Assembly of SNARE complexes that mediate neurotransmitter release requires opening of a ‘closed’ conformation of UNC-64/syntaxin. Rescue of *unc-13/Munc13* phenotypes by overexpressed open UNC-64/syntaxin suggested a specific function of UNC-13/Munc13 in opening UNC-64/ syntaxin. Here, we revisit the effects of open *unc-64*/syntaxin by generating knockin (KI) worms. The KI animals exhibited enhanced spontaneous and evoked exocytosis compared to wild-type animals. Unexpectedly, the open syntaxin KI partially suppressed exocytosis defects of various mutants, including *snt-1*/synaptotagmin, *unc-2/*P/Q/N-type Ca^2+^ channel alpha-subunit, and *unc-31/*CAPS in addition to *unc-13/*Munc13 and *unc-10/*RIM, and enhanced exocytosis in *tom-1/*Tomosyn mutants. However, open syntaxin aggravated the defects of *unc-18/*Munc18 mutants. Correspondingly, open syntaxin partially bypasses the requirement of Munc13 but not Munc18 for liposome fusion. Our results show that facilitating opening of syntaxin enhances exocytosis in a wide range of genetic backgrounds, and may provide a general means to enhance synaptic transmission in normal and disease states.

## Introduction

Synaptic vesicle exocytosis provides the chemical basis for neuronal communication, enabling the amazing diversity of functions of the brain. The SNARE (<u>s</u>oluble <u>N</u>SF-<u>a</u>ttachment <u>re</u>ceptor) proteins syntaxin, synaptobrevin/VAMP and SNAP-25 play a central role in exocytosis by forming a tight SNARE complex ^1^ that consists of a four-helix bundle ^2, 3^. This four-helix bundle is believed to be partially formed before Ca^2+^ influx [reviewed in ^4^]. Ca^2+^, acting through synaptotagmin-1 ^5, 6^, triggers full zippering of the SNARE complex, and ultimately results in fusion of the vesicle with the plasma membrane.

A biochemical step preceding the full fusion is often referred to as “priming”. It involves formation of the fusion-competent state of synaptic vesicles just prior to Ca^2+^ entry. The priming process is tightly controlled by multiple proteins, including Munc18-1/UNC-18 ^7–9^, Munc13-1/2/UNC-13 ^10–12^, Tomosyn ^13–15^ and RIM/UNC-10 ^16, 17^. However, the precise mechanisms through which these proteins mediate priming is still unclear.

Syntaxin/UNC-64 is a key SNARE protein that is crucial for exocytosis. It exists in an open and a closed conformation, the latter forming a tight complex with Munc18-1/UNC-18 ^18^. The binding of closed syntaxin and Munc18-1/UNC-18 has been shown to be critical for the level and plasmalemmal localization of syntaxin ^7, 19–22^. However, when held closed by Munc18-1/UNC-18, syntaxin is unable to interact with synaptobrevin and SNAP-25 to form SNARE complexes ^23, 24^. Munc13-1/UNC-13 was postulated to open syntaxin and consequently promote SNARE complex assembly and subsequent exocytosis at axon terminals. This important hypothesis originated from studies in which multi-copy over-expression of a constitutively open form of syntaxin/UNC-64 in *C. elegans* partially restored motility and strongly rescued synaptic vesicle exocytosis defects of *unc-13* null worms ^25^. The rescue was suggested to be due to a specific genetic interaction between *unc-13* and syntaxin, as expression of the open syntaxin in *unc-64* null (*js21*) mutant ^26^ did not enhance exocytosis compared with the wildtype syntaxin and did not suppress exocytosis defects of the P/Q/N-type calcium channel (UNC-2) mutant ^25^. This model also received support from diverse biochemical experiments, including the demonstration that the MUN domain of Munc13-1 enhances SNARE complex assembly starting from the Munc18-1-closed syntaxin-1 complex ^27–29^.

However, multicopy expression of open syntaxin was also found to rescue the exocytosis defects in *unc-10/*RIM mutants ^17^, and dense core vesicle docking defects in *unc-31/*CAPS mutants ^30^, calling into question the specificity of the genetic interaction between open syntaxin and *unc-13* as well as others. Strong overexpression of syntaxin may not inform on the full physiological state or functions of the endogenous gene products. Indeed, the extent of open syntaxin-mediated restoration of evoked neurotransmitter release in *unc-13* null mutant by these transgenes was variable in *C. elegans* ^15, 25^.

Moreover, the reported phenotype of multicopy open syntaxin in *C. elegans* was quite different from that of open syntaxin-1B knockin mice ^31, 32^. In particular, open syntaxin-1B led to enhanced Ca^2+^ sensitivity and fusion competency, as well as an increase in spontaneous release in hippocampal neurons ^31^ that was not observed in open syntaxin worms ^25^. In measurements performed at the Calyx of Held, the open syntaxin-1B mutation enhanced the speed of evoked release and accelerated fusion pore expansion ^32^. These phenotypes were postulated to arise from an increase in the number of SNARE complexes formed on vesicles, as a consequence of the facilitation of syntaxin-1B opening. However, use-dependent depression of EPSCs was increased in hippocampal synapses, suggesting a reduction in readily releasable pool (RRP) size in the open syntaxin-1B mutants due to reduced interactions between open syntaxin-1B and Munc18-1, and a corresponding decrease in the levels of both proteins ^33^. Furthermore, open syntaxin-1 was reported to fail or yield very limited rescue of phenotypes observed in Munc13-1 KO or Munc13-1/2 DKO mice ^31, 34^.

In this study, we re-evaluated the effects of the open form syntaxin by generating the first *in vivo* knock-in (KI) model of the open syntaxin mutant (L166A/E167A, often called LE mutant) in *C. elegans*. We then introduced the LE open mutation into various exocytosis mutant backgrounds to systematically examine the genetic interactions between open syntaxin and other regulators of exocytosis. The phenotype of open syntaxin and its genetic interactions with other synaptic transmission mutants only shared limited similarity with the ones previously described using multicopy transgenes ^25^. Our data show that the open syntaxin KI enhances neurotransmitter release and can partially rescue the phenotypes of a wide variety of exocytosis mutants except for Munc18/UNC-18. These results support the proposal that increasing the number of assembled SNARE complexes leads to enhanced neurotransmitter release and suggest that controlling the opening of syntaxin provides a general avenue to modulate synaptic transmission, which may help to develop novel therapeutic strategies for neurological diseases with impaired synaptic function.

## Results

### The open syntaxin knock-in (KI) mutant enhances spontaneous exocytosis compared to wild-type C. elegans

The L165A/E166A mutation in mammalian syntaxin-1A is known to result in a constitutively open form of syntaxin ^18^. Previous studies demonstrated that multicopy expression of UNC-64, the *C. elegans* syntaxin, harboring the analogous L166A and E167A mutations led to slower-moving animals than the wild-type without affecting the level of exocytosis ^25^. To determine the effects of a KI open syntaxin mutation on exocytosis, we generated *unc-64(sks4)* mutants where we introduced an L166A and E167A mutation in the *unc-64* endogenous locus by CRISPR/Cas9-directed homologous recombination (Fig. S1) ^35^. The genomic DNA sequence of open *syntaxin*/*unc-64* is shown in comparison with wild-type in Fig. S2. The corresponding Western blot analyses indicated that the open syntaxin/UNC-64 was expressed at slightly lower levels compared to wild-type UNC-64 expression (Fig. S1c). This reduced level of expression was consistent with a previous report on the knock-in model of open syntaxin-1B in mice ^33^, which is likely due to the lower affinity of open syntaxin binding for UNC-18 or Munc18-1 ^18, 36^, as the binary interaction between syntaxin and UNC-18/Munc18-1 seems important for the stability of syntaxin and Munc18-1 ^31^.

To investigate the effects of the KI, we first assayed open syntaxin mutants for their motility and aldicarb sensitivity (Fig. 1). Aldicarb is an acetylcholinesterase inhibitor and has been widely used to examine the level of acetylcholine release at the *C. elegans* neuromuscular junction ^37^. After 2 minutes in a liquid medium (M9 buffer), open syntaxin KI worms exhibited comparable (*p >* 0.05) thrashing rates to N2 wild-type worms (Fig. 1a). However, open syntaxin KI worms displayed a significantly faster onset of paralysis in the presence of aldicarb compared to N2 wild-type worms (Fig. 1b). To further examine whether open syntaxin increases synaptic vesicle release, we recorded miniature postsynaptic currents (mPSCs) at the *C. elegans* neuromuscular junction. We found a significantly increased frequency of mPSCs in the open syntaxin worms while we did not observe statistically significant changes in the amplitude of mPSCs (Fig. 1c, d). Our results indicate that, compared to wild-type syntaxin, the KI open syntaxin mutation significantly enhances spontaneous exocytosis. Thus, different from the results obtained from multicopy expression of open syntaxin in *unc-64* null animals ^25^, the phenotype of endogenously-expressed syntaxin KI animals was more consistent with the phenotype of syntaxin-1B knockin mice ^31, 32^.

**Figure 1.**
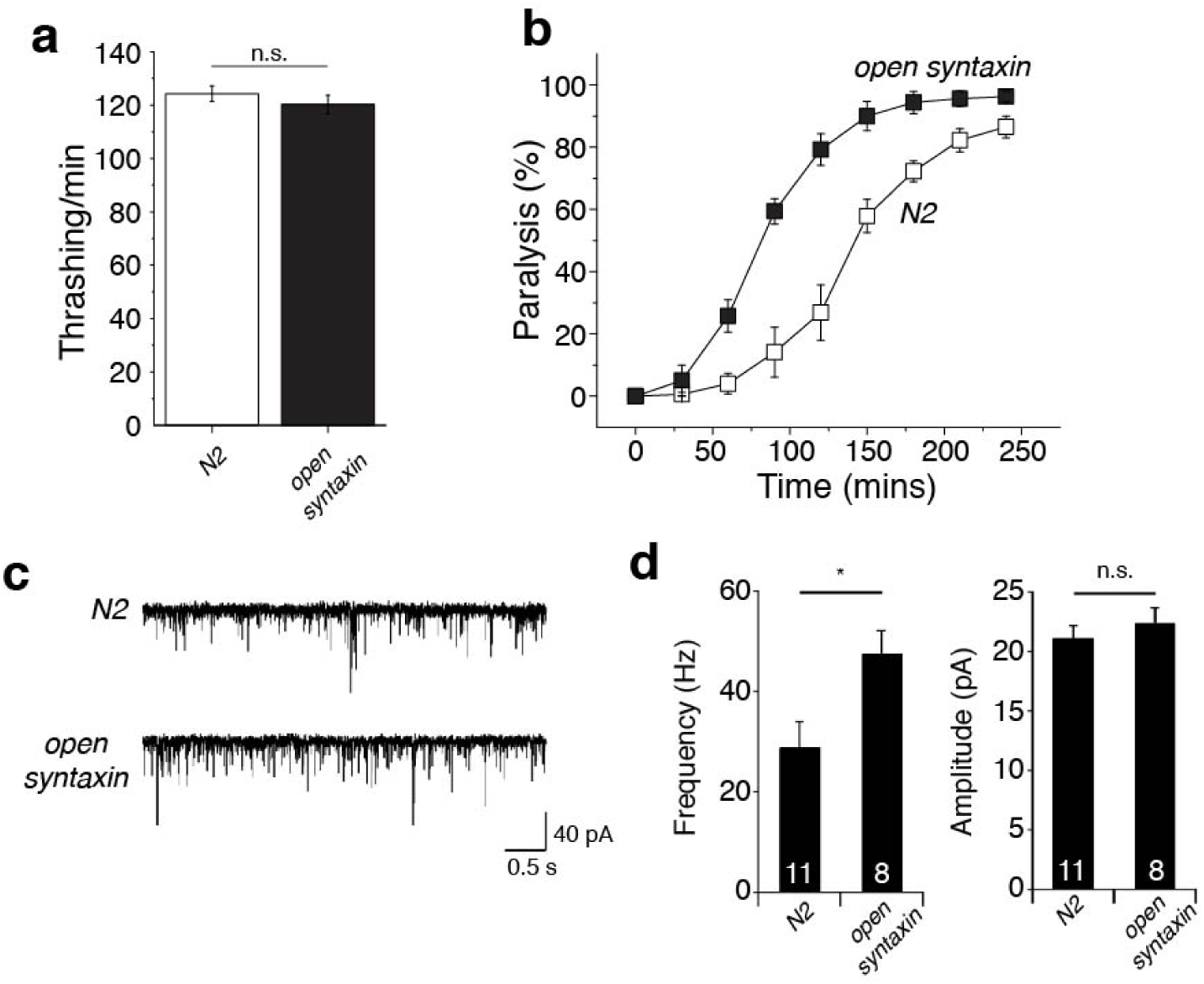
The knock-in open syntaxin mutation enhances exocytosis on a wild-type background. (a) Motility of each strain is measured by counting the number of thrashes per min in M9 buffer. Knock-in open syntaxin *C. elegans* display similar thrashing rates as wild-type *C. elegans* (122 thrashes/min vs 125 thrashes/min respectively). n=51 for each strain. In two-sample *t* tests, *t*_(100)_ = 0.745, *p =* 0.458. Error bars represent SEM. n.s. = not significant. (b) Aldicarb assays were conducted to measure aldicarb resistance of the animals. The *open syntaxin KI* mutant exhibits greater sensitivity to aldicarb compared to N2 wild-type worms. n=6. Each assay was conducted with 15-20 worms. Error bars represent SEM. (c) Representative traces of mPSCs recorded from N2 (top) and *open syntaxin KI* (bottom) worms. (d) Summary data of mPSC frequency (left) and amplitude (right) of N2 and *open syntaxin KI* worms. \**p* < 0.05; n.s. = not significant. n=11 for N2 and n=8 for *open syntaxin* animals.

### Open syntaxin partially rescues exocytosis defects in synaptotagmin-1 severe loss-of function mutants

Together with previous results showing that multicopy expression of open syntaxin-1 partially rescued the exocytosis defects of *unc-13* and *unc-10* mutants ^17, 25^, the increased spontaneous exocytosis caused by the KI open syntaxin mutation (Fig. 1) suggested the possibility that the open syntaxin mutation may provide a general means to enhance exocytosis, and thus overcome defects caused by diverse types of mutations in the exocytotic machinery.

In this context, the ability of open syntaxin to rescue a null mutant of synaptotagmin-1/*snt-1*, a key calcium sensor for exocytosis ^5, 6^, has not been investigated. We crossed a severe *snt-1* loss-of-function mutant allele, *snt-1(md290)* ^38^, with the open syntaxin KI mutant to assay for thrashing ability and aldicarb sensitivity (Fig. 2). Importantly, we made the novel finding that open syntaxin can partially rescue synaptotagmin-1 mutant phenotypes in *C. elegans.* Thus, our *open syntaxin KI*; *snt-1(md290)* double mutants exhibited greater thrashing rates and paralysis in response to aldicarb than those observed for the *snt-1(md290)* mutant (Fig. 2a, 2b). Analysis of the mPSCs showed that open syntaxin increased the frequency of mPSCs but not the amplitude when compared to *snt-1* mutants alone (Fig. 2c, 2d). These results suggest that open syntaxin can bypass the requirement of a wider range of exocytosis regulator than previously recognized, including synaptotagmin.

**Figure 2.**
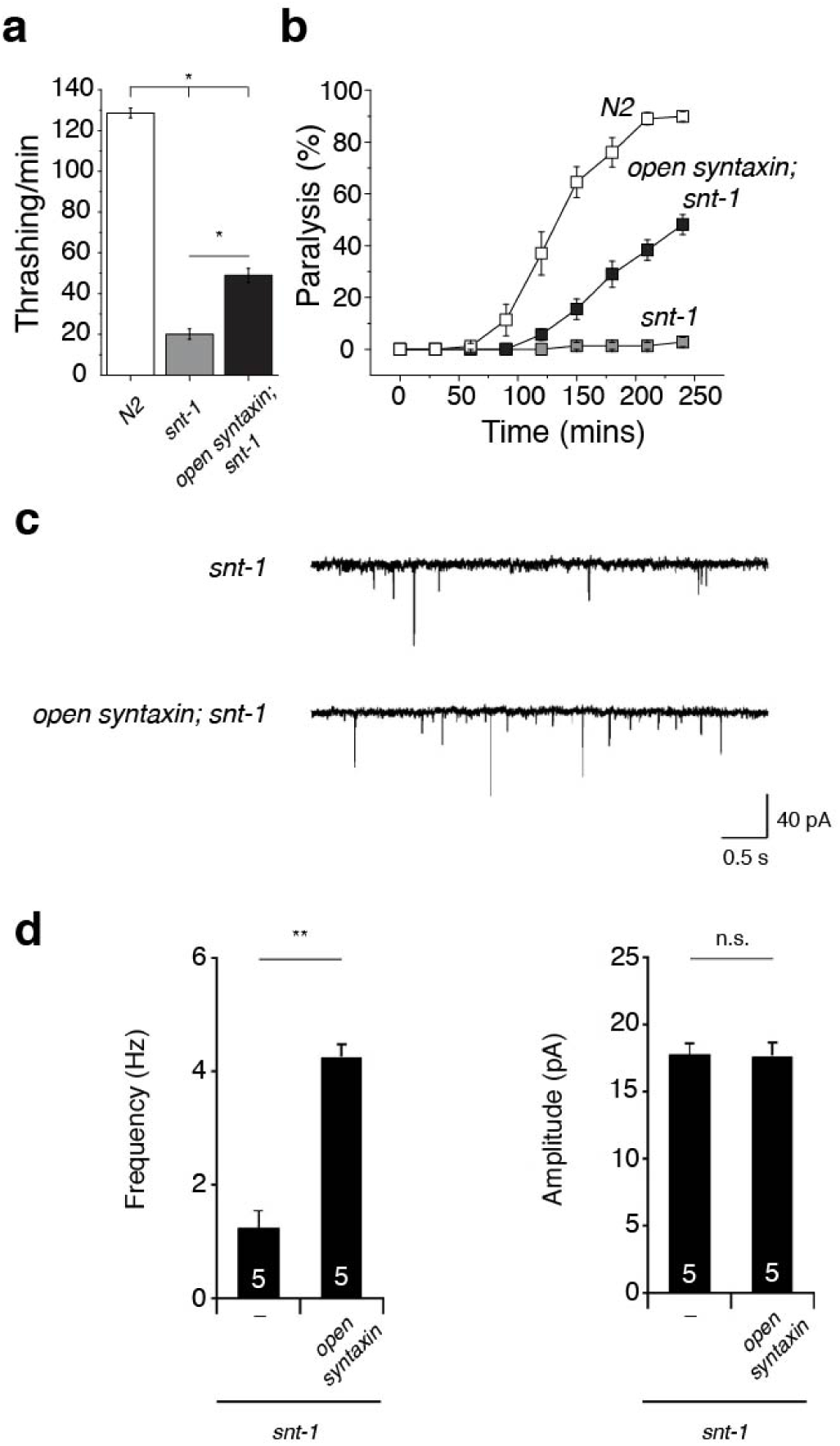
Open syntaxin rescues acetylcholine secretion defects in *snt-1* null *C. elegans*. (a) Thrashing assay of N2, *snt-1,* and *open syntaxin; snt-1* double mutants in M9 buffer. *snt-1* worms displayed impaired thrashing rates (20.1 thrashes/min) which was increased by the introduction of open syntaxin in the double mutant (48.9 thrashes/min). n=40 for each strain. In one-way ANOVA statistical tests, *F*_(2,117)_ = 364 and *p =* 0. Tukey’s test was performed for means analysis in ANOVA. Error bars represent SEM. \**p* < 0.05. (b) Aldicarb assays of N2, *snt-1,* and *open syntaxin; snt-1* double mutants. *open syntaxin; snt-1* animals displayed slightly increased sensitivity to aldicarb compared to *snt-1* single mutants which were resistant to aldicarb’s paralyzing effects. n=6. Each assay was conducted with 15-20 worms. Error bars represent SEM. (c) Representative mPSC traces recorded from *snt-1,* and *open syntaxin; snt-1* worms. (d) Summary data of mPSC frequency (left) and amplitude (right) of *snt-1* and *open syntaxin; snt-1* worms. \*\**p* < 0.01; n.s. = not significant. n=5 for *snt-1(md290)* and *open syntaxin; snt-1(md290)* animals.

### The open syntaxin knock-in (KI) mutant enhances evoked neurotransmitter release compared with wild-type syntaxin and partially restores the defects of evoked release of snt-1 mutants

In *C. elegans* neuromuscular junctions, reliable recording of evoked neurotransmitter release by electrical stimulation has been a challenge. To overcome this technical difficulty, optogenetic stimulation has been developed in which Channelrhodopsin-2 is expressed specifically in cholinergic (*zxls6* strain) neurons and evoked release is triggered by light ^39^.

To examine the effects of open syntaxin on evoked acetylcholine release, we generated *open syntaxin KI* as well as *open syntaxin KI; snt-1(md290)* mutants in the *zxls6* background. When stimulated by 10-ms blue light (3.75 mW/mm^2^), open *syntaxin KI* mutants exhibited a significant increase in the half-width and charge transfer of evoked excitatory postsynaptic currents (EPSC) compared to wild-type (Fig. 3a, d, e). However, the peak amplitude was unchanged (Fig. 3b). *Open syntaxin KI; snt-1(md290)* mutants exhibited increases in the amplitude as well as the half-width and charge transfer of EPSCs compared with *snt-1(md290)* animals (Fig. 3a, b, d, e). The rise time to the peak amplitude (10-90%) is unchanged in all the strains tested (Fig. 3c). Taken together, our results show that open syntaxin enhances not only the frequency of the spontaneous exocytosis (Figs. 1c, d, 2c, d) but also the amount of acetylcholine released by light-induced depolarization. Furthermore, it partially rescues the defects of evoked exocytosis of *snt-1* mutant animals.

**Figure 3.**
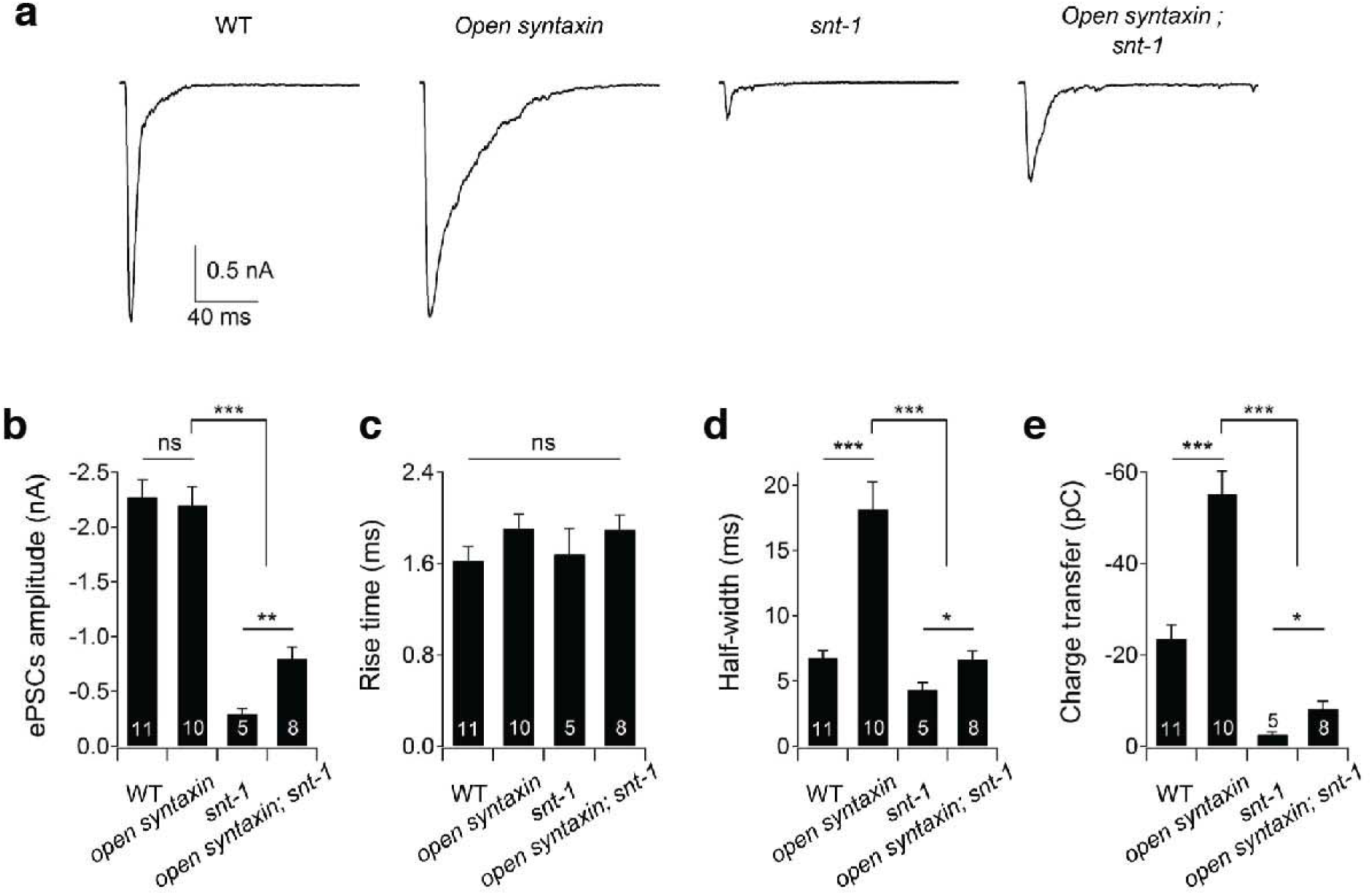
The knock-in open syntaxin mutation enhances evoked neurotransmitter release compared with the wild-type and partially restores the defects of evoked release of *snt-1* mutants. (a) Representative EPSCs traces recorded from wild-type, *open syntaxin, snt-1,* and *open syntaxin; snt-1* worms in the *zxIs6* transgenic background with 10 ms blue light illumination (3.75 mW/mm^2^). Muscle cells were recorded with a holding potential of 60 mV.L (b-e) Summary data of EPSCs amplitude (b), rise time (10-90%) (c), half-width (d) and total charge transfer (e) of wild-type, *open syntaxin, snt-1,* and *open syntaxin; snt-1* worms, respectively. \**p* < 0.05, \*\**p* < 0.01, \*\*\**p* < 0.001; n.s. = not significant. Error bars represent SEM.

### Open syntaxin rescues exocytosis in unc-2 mutant C. elegans

The key evidence that supported a specific genetic interaction between open syntaxin and UNC-13 derived from the observation that open syntaxin did not rescue exocytosis of the *unc-64*(*js21*) null mutant better than the wild-type syntaxin and did not overcome the impaired synaptic exocytosis of a severe loss-of-function mutant (*e55*) of *unc-2* (Richmond et al., 2001). UNC-2 is an alpha subunit of P/Q/N-type Ca_V_2-like voltage gated calcium channels, which control the influx of calcium upstream of synaptic vesicle exocytosis ^40^. The *e55* mutant has a Q571Stop mutation in the Ca_V_2-like voltage gated calcium channel alpha subunit, which reduces calcium current and leads to impaired locomotion and exocytosis ^40, 41^. Since our *open syntaxin KI*, or *unc-64(sks4)*, increases synaptic exocytosis compared to wild-type N2 (Fig. 1, 3) and partially rescues *snt-1* phenotypes (Fig. 2, 3), we hypothesized that it might also rescue the reduced exocytosis of the *unc-2* loss-of-function mutant (*e55*).

*Unc-2(e55)* worms exhibited reduced thrashing and aldicarb sensitivity, whereas *open syntaxin KI; unc-2(e55)* double mutants displayed robustly rescued thrashing (*p <* 0.01) compared to the *unc-2(e55)* mutant alone, and an aldicarb sensitivity comparable to the wild-type level (Fig. 4A, B). Analysis of the mPSCs showed that wild-type (N2), *unc-2(e55),* and the open syntaxin; *unc-2(e55)* double mutants displayed similar amplitudes, but there was markedly reduced mPSC frequency in the *unc-2(e55)* worms that were rescued by *open syntaxin KI* in the double mutant (Fig. 4c, d). These striking results contrast with those obtained with multicopy expression of open syntaxin-1 ^25^, and provide compelling evidence that the open syntaxin mutation can at least partially overcome widely diverse defects in exocytosis.

**Figure 4.**
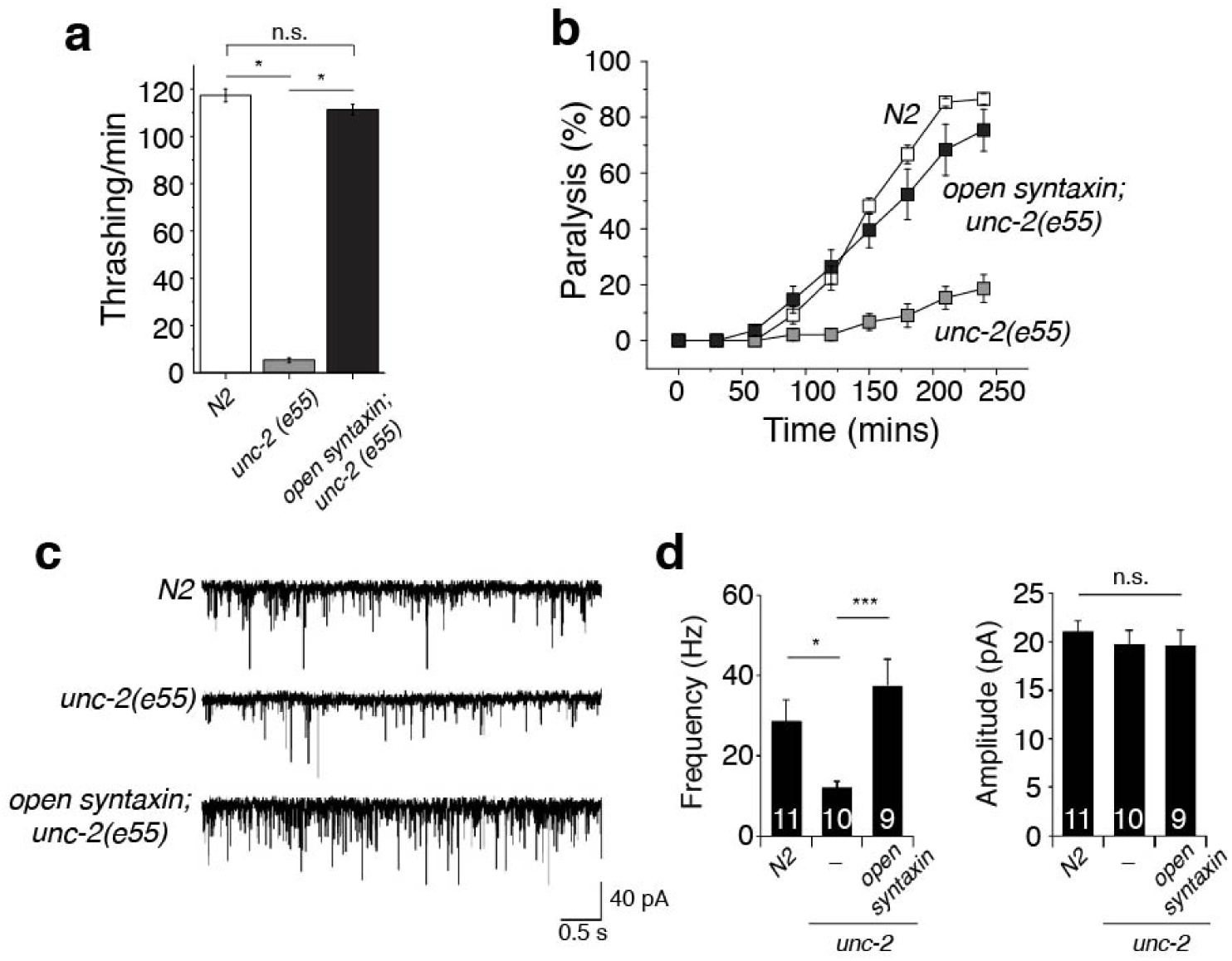
The knock-in open syntaxin mutation rescues the behavioral and exocytosis defects of *unc-2* null *C. elegans.* (a) Thrashing assay of N2, *unc-2(e55),* and *open syntaxin; unc-2(e55)* double mutants in M9 buffer. *unc-2(e55)* worms displayed greatly reduced thrashing rates (5.43 thrashes/min) which was rescued by the introduction of open syntaxin in the double mutant. n=40 for each strain. In one-way ANOVA statistical tests, *F*_(2,117)_ = 836 and *p =* 0. Tukey’s test was performed for means analysis in ANOVA. Error bars represent SEM. \**p* < 0.05; n.s. = not significant. (b) Aldicarb assays of N2, *unc-2(e55),* and *open syntaxin; unc-2(e55)* double mutants. *unc-2* animals displayed resistance to aldicarb (grey), while the double mutant restored aldicarb sensitivity to near wild-type levels (black). n=6. Each assay was conducted with 15-20 worms. Error bars represent SEM. (c) Representative mPSC traces recorded from N2, *unc-2(e55),* and *open syntaxin; unc-2(e55)* worms. (d) Summary data of mPSC frequency (left) and amplitude (right) of N2, *unc-2(e55),* and *open syntaxin; unc-2(e55)* worms. \**p* < 0.05; *** *p* < 0.001; n.s. = not significant. n=11 for N2, n=10 for *unc-2(e55)* and n=9 for *open syntaxin; unc-2(e55)* animals.

We also tested the effect of open syntaxin on a gain-of-function allele of *unc-2*(*hp647*) (Fig. S3). The *hp647* allele contains a L653F mutation ^42^. A gain-of-function mutant allele of *unc-2, unc-2(hp647)* exhibited striking hypersensitivity to aldicarb (Fig. S3b), suggesting that this mutant increased acetylcholine release. Interestingly however, thrashing frequency in *unc-2(hp647)* was not greater than that of the N2 wild-type (Fig. S3a). *Open syntaxin KI; unc-2(hp647)* double mutants appeared to exhibit a slight further increase of aldicarb sensitivity when compared with *unc-2(hp647)* alone (Fig. S3b), but it was difficult to manifest this effect using 1 mM aldicarb concentration. To further elucidate the difference between the strains, we used a lower aldicarb concentration plate (0.3 mM) and monitored for paralysis every 15 minutes (Supplementary Fig. S3c). We observed that *open syntaxin KI* slightly enhanced the aldicarb sensitivity of the *unc-2* gain-of-function mutant. Thus, open syntaxin can enhance exocytosis in both loss-of-function and gain-of-function mutants of the calcium channel UNC-2. These results support the possibility that open syntaxin increases synaptic transmission regardless of genetic background.

### Open syntaxin weakly enhances exocytosis in unc-13 mutants

Unlike multicopy expression of open syntaxin in *unc-64* null background, our KI open syntaxin enhanced exocytosis compared to the N2 wild-type (Fig. 1) and rescued exocytosis in *unc-2* loss-of-function mutant (Fig. 4). Therefore, we further investigated whether rescue of *unc-13* null *C. elegans* phenotypes by our open syntaxin KI mutation might differ from the results obtained with multicopy expression of open syntaxin ^15, 25^. We found that the severe thrashing defect of *unc-13(s69)* mutant (0.0125/min) was slightly alleviated by open syntaxin KI, which increased thrashing of *open syntaxin KI; unc-13(s69)* double mutant to 0.713/min (Fig. 5a). However, this effect was not as strong as the effect of multicopy open syntaxin, which was reported to be ∼6/min ^25^. Moreover, we did not observe a significant increase in aldicarb sensitivity when compared to the *unc-13(s69)* mutant (*p* > 0.05) after the usual 4-hour exposure (Fig. 5b). After 24 hours in the aldicarb plate, approximately 33% of *open syntaxin KI; unc-13(s69)* worms were paralyzed, compared to the 7% paralyzed *unc-13(s69)* worms. All wild-type worms paralyzed at the end of the 24 hours. Thus, there is only a weak rescue of acetylcholine release from *unc-13(s69)* animals.

**Figure 5.**
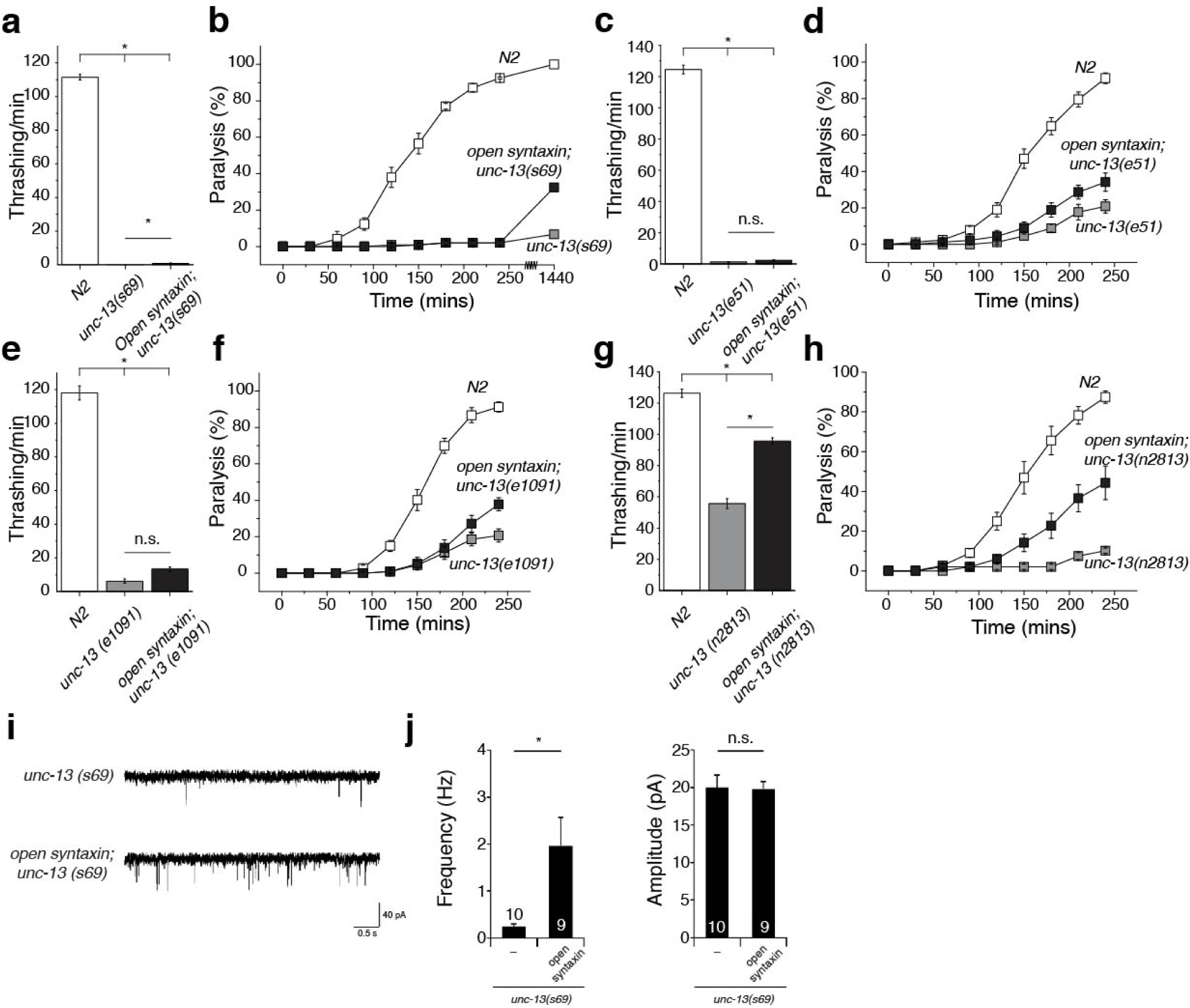
Open syntaxin weakly rescues the behavioral and exocytosis defects of *unc-13* mutants. (a) Thrashing assay of N2, *unc-13(s69),* and *open syntaxin; unc-13(s69)* double mutants in M9 buffer. *unc-13(s69)* worms displayed greatly reduced thrashing rates (0.0125 thrashes/min) which was slightly increased by the introduction of open syntaxin in the double mutant (0.713 thrashes/min). n=40 for each strain. In one-way ANOVA statistical tests, *F*_(2,117)_ = 4270 and *p =* 0. Tukey’s test was performed for means analysis in ANOVA. Error bars represent SEM. \**p* < 0.05. (b) Aldicarb assays of N2, *unc-13(s69),* and *open syntaxin; unc-13(s69)* double mutants. *unc-13(s69)* and *open syntaxin; unc-13(s69)* animals both displayed resistance to aldicarb. After 24 hours on aldicarb, 33% of *open syntaxin; unc-13(s69)* worms were paralyzed, whereas 7% of *unc-13(s69)* worms were paralyzed. n=6 for 4-hr assays, n=1 for 24-hr assay. Each assay was conducted with 15-20 worms. Error bars represent SEM. (c) Thrashing assay of N2, *unc-13(e51),* and *open syntaxin; unc-13(e51)* double mutants in M9 buffer. *unc-13(e51)* worms displayed greatly reduced thrashing rates (1.31 thrashes/min) which was slightly (but not significantly) increased by the introduction of open syntaxin in the double mutant (2.31 thrashes/min). n=57 for each strain. In one-way ANOVA statistical tests, *F*_(2,168)_ = 1760 and *p =* 0.37. Tukey’s test was performed for means analysis in ANOVA. Error bars represent SEM. \**p* < 0.05. (d) Aldicarb assays of N2, *unc-13(e51),* and *open syntaxin; unc-13(e51)* double mutants. *open syntaxin; unc-13(e51)* animals displayed slightly increased sensitivity to aldicarb compared to *unc13(e51)* single mutants. n=6. Each assay was conducted with 15-20 worms. Error bars represent SEM. (e) Thrashing assay of N2, *unc-13(e1091),* and *open syntaxin; unc-13(e1091)* double mutants in M9 buffer. *unc-13(e1091)* worms displayed greatly reduced thrashing rates (6.14 thrashes/min) which was moderately (but not significantly) increased by the introduction of open syntaxin in the double mutant (13.3 thrashes/min). n=40 for each strain. In one-way ANOVA statistical tests, *F*_(2,117)_ = 574 and *p =* 0.13. Tukey’s test was performed for means analysis in ANOVA. Error bars represent SEM. \**p* < 0.05. (f) Aldicarb assays of N2, *unc-13(e1091),* and *open syntaxin; unc-13(e1091)* double mutants. *open syntaxin; unc-13(e1091)* animals displayed slightly increased sensitivity to aldicarb compared to *unc13(e1091)* single mutants. n=6. Each assay was conducted with 15-20 worms. Error bars represent SEM. (g) Thrashing assay of N2, *unc-13(n2813),* and *open syntaxin; unc-13(n2813)* double mutants in M9 buffer. *unc-13(n2813)* worms displayed reduced thrashing rates (55.7 thrashes/min) which was increased by the introduction of open syntaxin in the double mutant (95.6 thrashes/min). n=40 for each strain. In one-way ANOVA statistical tests, *F*_(2,117)_ = 173 and *p =* 0. Tukey’s test was performed for means analysis in ANOVA. Error bars represent SEM. \**p* < 0.05. (h) Aldicarb assays of N2, *unc-13(n2813),* and *open syntaxin; unc-13(n2813)* double mutants. *open syntaxin; unc-13(n2813)* animals displayed significantly increased sensitivity to aldicarb compared to *unc13(n2813)* single mutants. n=6. Each assay was conducted with 15-20 worms. Error bars represent SEM. (i) Representative mPSC traces recorded from N2, *unc-13(s69),* and *open syntaxin; unc-13(s69)* worms. (j) Summary data of mPSC frequency (left) and amplitude (right) of N2, *unc-13(s69),* and *open syntaxin; unc-13(s69)* worms. \**p* < 0.05; \*\**p* < 0.01; n.s. = not significant. n=10 for *unc-13(s69)* and n=9 for *open syntaxin; unc-13(s69)* animals.

Previous studies using electrophysiological analysis reported conflicting levels of the rescue of synaptic vesicle exocytosis in *unc-13(s69)* animals by multicopy expression of open syntaxin ^15, 25^. We therefore performed electrophysiological analyses of the effects of *open syntaxin KI* on *unc-13(s69)* mutants. The amplitudes of the mPSCs in the *unc-13(s69)* mutant were similar to those of the N2 worms, but the frequency of mPSCs were drastically reduced (Fig. 5i, 5j; compare with Fig. 1c, 1d). The open syntaxin KI mutation on the *unc-13(s69)* background enhanced mPSC frequently significantly from <0.5 Hz to ∼2 Hz (Fig. 5j).

Nevertheless, the enhanced level did not reach wild-type levels (∼30-40 Hz; Fig. 1d). The weak rescue phenotype in electrophysiology is more consistent with the weak rescue observed in *C. elegans* ^15^ and the phenotype of Munc13-1/2 DKO neurons with overexpressed open syntaxin-1^34^.

The UNC-13 protein exists as LR and MR isoforms ^12, 43^. *unc-13(s69)* mutants possess a C1029fs mutation in both isoforms that results in a severe deleterious phenotype such that rescue with open syntaxin may be minimal. Therefore, we investigated whether the KI open syntaxin can rescue weaker loss-of-function alleles of *unc-13*: *e51*, *e1091*, and *n2813*. *e51* and *e1091* mutants are null in their LR isoform only, while *n2813* mutants possess a partial loss of function gene that results in a single S1574R missense mutation in both UNC-13 isoforms ^12^. *Open syntaxin KI*; *unc-13(e51)* double mutants did not show rescued thrashing (*p* = 0.37) but did display increased sensitivity to aldicarb compared to the *unc-13(e51)* mutant alone (Fig. 5c, 5d). Similarly, *open syntaxin KI*; *unc-13(e1091)* double mutants did not show rescued thrashing (*p* = 0.13) but did exhibit increased sensitivity to aldicarb compared to the *unc-13(e1091)* alone (Fig. 5e, 5f). Finally, the *open syntaxin KI*; *unc-13(n2813)* double mutant displayed rescue in thrashing (*p* < 0.05) compared to the *unc-13(n2813)* mutant alone (Fig. 5g). Aldicarb sensitivity in the *open syntaxin KI*; *unc-13(n2813)* double mutant was also enhanced compared to the *unc-13(n2813)* mutant alone (Fig. 5h). Overall, our results suggest that open syntaxin increases acetylcholine release from *unc-13* mutants, but the level of rescue is rather limited and is less pronounced than the rescue of the *unc-2* loss-of-function mutant.

### Open syntaxin and unfurled unc-18 differentially interact with tom-1 in the rescue of unc-13 null mutant

A limited rescue of *unc-13(s69)* by open syntaxin KI could be due to too severe impairment of synaptic vesicle priming in the absence of *unc-13*. In this context, the *tom-1* mutant that lacks Tomosyn, a protein that competes with synaptobrevin for SNARE complex formation ^44^, partially rescued *unc-13* phenotypes, and multicopy expression of the open syntaxin mutant further increased the rescue to a modest extent ^15^. In contrast, we recently discovered an interesting synergism between a KI of an *unc-18* mutant containing a point mutation (P334A) that is believed to unfurl a loop involved in synaptobrevin binding and the *tom-1(ok285)* null mutant in rescuing the *unc-13(s69)* mutant in synaptic vesicle exocytosis ^45^. That is, although *unc-18(P334A) KI* or *tom-1(ok285)* had a limited rescuing effect in aldicarb sensitivity of the *unc-13(s69)* mutant, in combination they restored aldicarb sensitivity to the wild-type level. Therefore, we investigated whether introducing open syntaxin as a KI can restore *unc-13(s69)* strongly in the *tom-1* null background.

We first examined whether the effects of open syntaxin and *tom-1* on acetylcholine release and thrashing. As previously reported ^15, 45^, *tom-1* null exhibited slightly reduced motility and thrashing compared to N2 wild-type despite the increased acetylcholine release (Fig. 6a, b). Importantly, although open syntaxin KI did not affect the level of thrashing on *tom-1(ok285)* mutant, it further increased aldicarb sensitivity (Fig. 6b). These additive effects on aldicarb sensitivity suggest that facilitation of SNARE complex assembly by open syntaxin and the absence of TOM-1 can cooperate to further enhance exocytosis. In the triple mutants of *open syntaxin KI*; *tom-1 null; unc-13(s69),* we observed that the open syntaxin partially increased motility and acetylcholine release of *unc-13* in the absence of *tom-1* (Fig. 6c, d). However, the rescue measured by the sensitivity to aldicarb is not as strong as that caused by the unfurled *unc-18(sks2)* mutant in the absence of TOM-1 and UNC-13. As we previously reported, the aldicarb sensitivity of unfurled *unc-18(sks2) KI*; *tom-1(ok285)*; *unc-13(s69)* reached N2 wild-type levels (Fig. 6f) ^45^. These results suggest that open syntaxin and *tom-1 null* exhibit additive, not synergistic effects on the rescue of *unc-13(s69)*. We conclude that the ability of open syntaxin to suppress *unc-13* null is limited even in the absence of TOM-1.

**Figure 6.**
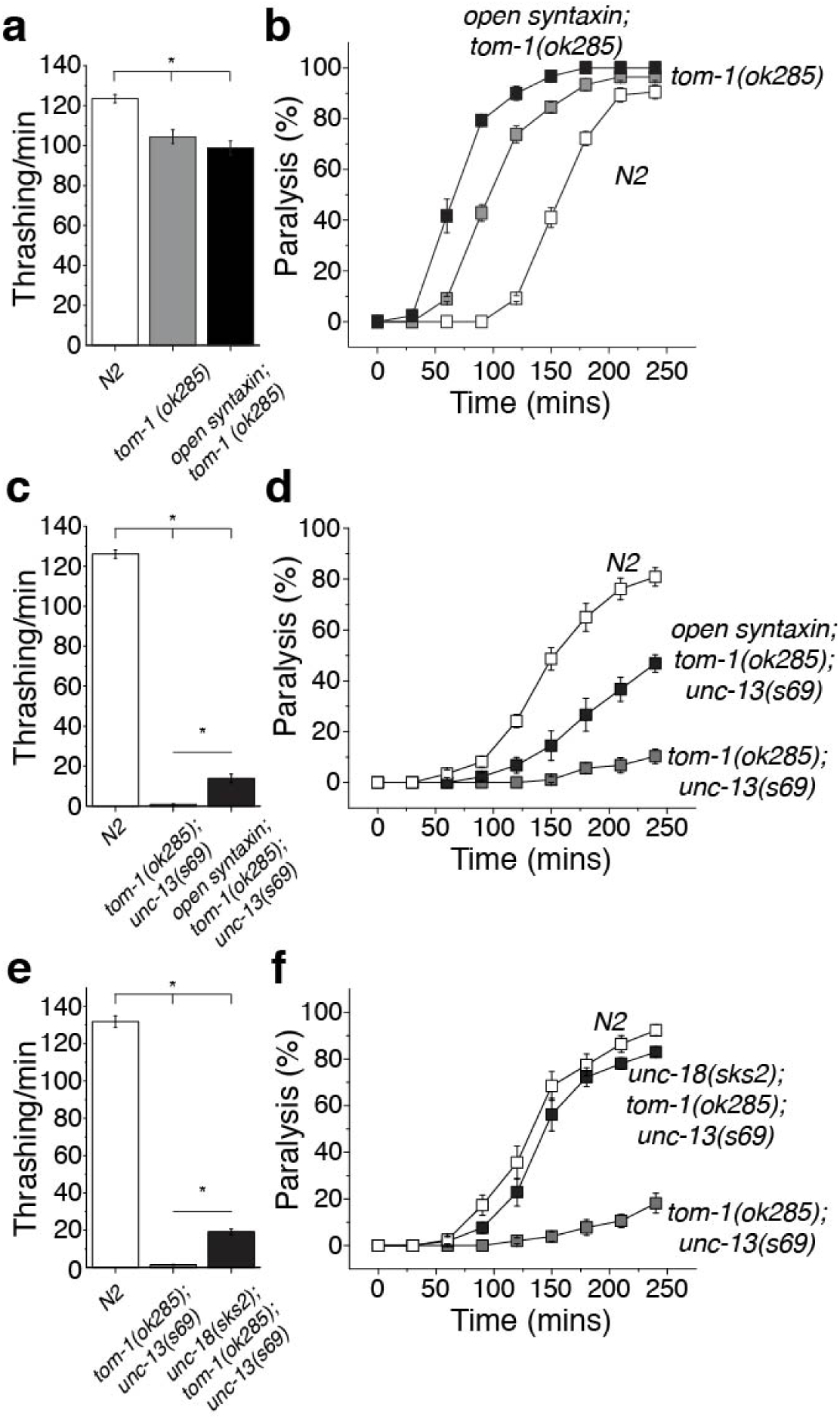
Open syntaxin and unfurled *unc-18* differentially interact with *tom-1* in the rescue of *unc-13.* (a) Thrashing assay of N2, *tom-1(ok285)* and *open syntaxin; tom-1* double mutants in M9 buffer. *tom-1* and *open syntaxin; tom-1* thrashing rates were comparable to each other (104 thrashes/min vs 98.9 thrashes/min respectively) while moderately lower than the N2 wild-type (123.5 thrashes/min). n=40 for each strain. In one-way ANOVA statistical tests, *F*_(2,117)_ = 16.9 and *p =* 0. Tukey’s test was performed for means analysis in ANOVA. Error bars represent SEM. \**p* < 0.05; n.s. = not significant. (b) Aldicarb assays of N2, *tom-1(ok285)* and *open syntaxin; tom-1* double mutants. *open syntaxin; tom-1* animals displayed significantly increased sensitivity to aldicarb compared to *tom-1* single mutants, while both displayed hypersensitivity to aldicarb compared to N2. n=6. Each assay was conducted with 15-20 worms. Error bars represent SEM. (c) Thrashing assay of N2, *tom-1; unc-13(s69),* and *open syntaxin; tom-1; unc-13(s69)* mutants in M9 buffer. The triple mutant significantly rescued thrashing compared to the *tom-1; unc-13(s69)* double mutant thrashing rate (13.9 thrashes/min vs 0.938 thrashes/min). n=40 for each strain. In one-way ANOVA statistical tests, *F*_(2,117)_ = 1640 and *p =* 0. Tukey’s test was performed for means analysis in ANOVA. Error bars represent SEM. \**p* < 0.05. (d) Aldicarb assays of N2, *tom-1; unc-13(s69),* and *open syntaxin; tom-1; unc-13(s69)* mutants. *open syntaxin; tom-1; unc-13(s69)* animals displayed significantly increased sensitivity to aldicarb compared to *tom-1; unc-13(s69)* double mutants. n=6. Each assay was conducted with 15-20 worms. Error bars represent SEM. (e) Thrashing assay of N2, *tom-1; unc-13(s69),* and *unc-18(sks2); tom-1; unc-13(s69)* mutants in M9 buffer. The triple mutant significantly rescued thrashing compared to the *tom-1; unc-13(s69)* double mutant thrashing rate (19.2 thrashes/min vs 1.48 thrashes/min respectively). n=40 for each strain. In one-way ANOVA statistical tests, *F*_(2,117)_ = 1210 and *p =* 0. Tukey’s test was performed for means analysis in ANOVA. Error bars represent SEM. \**p* < 0.05. (f) Aldicarb assays of N2, *tom-1; unc-13(s69),* and *unc-18(sks2); tom-1; unc-13(s69)* mutants. *unc-18(sks2); tom-1; unc-13(s69)* animals displayed significantly increased sensitivity to aldicarb compared to *tom-1; unc-13(s69)* double mutants. n=6. Each assay was conducted with 15-20 worms. Error bars represent SEM.

### Rescue of unc-10 and unc-31 by open syntaxin KI

Multicopy expression of open syntaxin-1, which can rescue the *unc-13(s69)* mutant, was also reported to rescue exocytosis defects of *unc-10* mutant ^17^ as well as docking defects of dense core vesicles of *unc-31* mutant. UNC-10 can bind and activate UNC-13 ^46–48^. UNC-31 is involved in dense core vesicle exocytosis but may also influence synaptic vesicle exocytosis ^49, 50^. To determine whether open syntaxin KI indeed enhances exocytosis of these synaptic exocytosis mutants, we crossed severe loss-of-function mutants, *unc-10(md1117)* and *unc-31(e928) C. elegans,* with open syntaxin KI mutants to generate double mutants, and assayed for thrashing ability and aldicarb sensitivity (Fig. S4). The *open syntaxin KI; unc-10(md1117)* double mutants displayed moderate rescue in thrashing rates (*p <* 0.05) as well as aldicarb sensitivity (Fig. S4a, S4b). Similarly, *open syntaxin KI; unc-31(e928)* double mutants also displayed moderate rescue of thrashing (*p <* 0.05), but restored aldicarb sensitivity to levels similar to those of wild-type worms (Fig. S4c, S4d). These results further show that open syntaxin can suppress exocytosis defects from a wide range of synaptic transmission mutants. Furthermore, the degree of rescue of these mutants seems to be comparable to the rescue of *unc-13* mutants and in some cases is more effective.

### Open syntaxin aggravates the phenotype of unc-18 loss-of-function mutants

Thus far, open syntaxin KI increased exocytosis to some degree in all the genetic backgrounds that we tested. Previous work suggested that multicopy over-expression of syntaxin does not bypass the requirement of UNC-18 ^9^; we thus sought to investigate whether open syntaxin KI can rescue *unc-18* null worms. In the present study, we tested 3 strains of *unc-18* severe loss-of-function mutants: *md299, e81, and sks1.* The *md299* strain possesses a multigenic deletion resulting in the loss of the promoter and open reading frame of the *unc-18* gene ^9^. The *e81* allele contains a C1582T mutation in exon 9, resulting in a Q530Stop mutation and is thus likely null of the *unc-18* gene ^51^. Finally, the *sks1* strain contains a large insertion containing the genes for selection markers GFP and neomycin that disrupts proper transcription of the *unc-18* gene ^45^.

In all three *unc-18* alleles, thrashing rate was severely reduced and very little to no aldicarb sensitivity was observed (Figs. 7a,b, S5a,b, S6a,b). Nevertheless, the *md299* strain exhibited measurable thrashing activity as previously reported ^45^. Interestingly, *open syntaxin KI; unc-18(md299)* strongly reduced this thrashing activity (Fig. 7a). Thus, KI open syntaxin aggravated the motility defects of *unc-18(md299)*. The double mutant and *md299* mutant displayed similarly poor aldicarb sensitivity. Therefore, worms were left overnight in the aldicarb plate and examined after 24 hours for paralysis. At the end of the 24-hour period, 100% of wild-type worms were paralyzed, while approximately 7% of the *unc-18(md299)* worms were paralyzed, and double mutants did not paralyze on 1 mM aldicarb even after 24 hours (Fig. 7b). The motility and aldicarb sensitivity of *e81* and *sks1* mutants were too low to detect a putative aggravating effect of open syntaxin KI (Fig. S5, S6).

**Figure 7.**
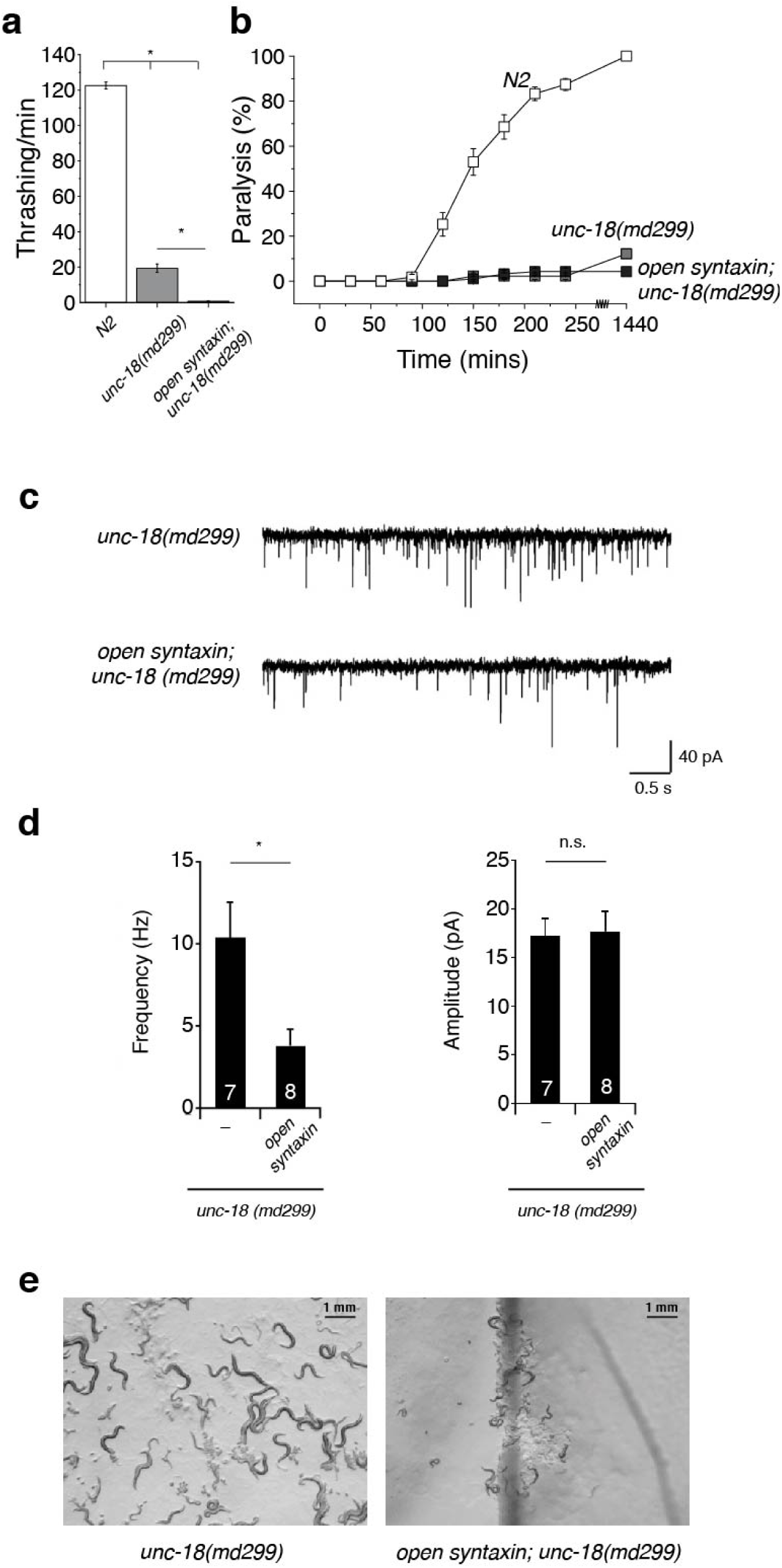
Open syntaxin further impairs the *unc-18* null phenotype. (a) Thrashing assay of N2, *unc-18(md299),* and *open syntaxin; unc-18(md299)* double mutants in M9 buffer. *unc-18(md299)* worms displayed greatly reduced thrashing rates (19.4 thrashes/min) which was reduced further by the introduction of open syntaxin in the double mutant to 0.610 thrashes/min. n=41 for each strain. In one-way ANOVA statistical tests, *F*_(2,120)_ = 1350 and *p =* 0. Tukey’s test was performed for means analysis in ANOVA. Error bars represent SEM. \**p* < 0.05. (b) Aldicarb assays of N2, *unc-18(md299),* and *open syntaxin; unc-18(md299)* double mutants. *unc-18* and *open syntaxin; unc-18* animals displayed similar resistance to aldicarb even after 24 hours. n=6 for 4-hr assays, n=1 for 24-hr assay. Each assay was conducted with 15-20 worms. Error bars represent SEM. (c) Representative mPSC traces recorded from *unc-18(md299)* and *open syntaxin; unc-18(md299)* worms. (d) Summary data of mPSC frequency (left) and amplitude (right) of *unc-18(md299)* and *open syntaxin; unc-18(md299)* worms. \**p* < 0.05; n.s. = not significant. n=7 for *unc-18(md299)* and n=8 for *open syntaxin; unc-18(md299)* animals. (e) Growth rate/body size images taken at day 9.

We examined the effects of open syntaxin KI on *unc-18* mutants using electrophysiology. *md299* mutants showed a reduced mPSC frequency compared to WT (compare Fig. 7c, 7d with Fig. 1c, 1d), and open syntaxin further reduced mPSC frequency in an *md299* background (Fig. 7c, 7d). The amplitude of mPSCs showed no difference between *md299* and *open syntaxin KI; md299* mutants. Despite our best efforts, *open syntaxin KI; unc-18*(*e81)* double mutants were too small and sick to perform reliable neuromuscular junction preparations for electrophysiology.

We indeed observed the striking effects of KI open syntaxin on the growth rate and body size of all the *unc-18* loss-of-function mutants. The slower growth rate and smaller size of *unc-18* single mutants were previously reported (Sassa et al., 1999). Starting from 15 eggs (day 0) in N2 wild-type worms, by day 7 the plate is populated with adults and the OP50 bacterial lawn is depleted. Because of their impaired movement, the various *unc-18* mutants were unable to fully populate the NGM plate and deplete the entire OP50 bacterial lawn. However, the areas of the plate where they were located were completely populated with smaller-sized adults and depleted of OP50 by day 9. In contrast, the *open syntaxin KI; unc-18* mutants failed to populate the plate or deplete the bacterial lawn to the same extent as the *unc-18* mutants by day 9 (Fig. 7e, Fig.S5c, S6c). Our results strongly suggest that unlike all other synaptic transmission mutants, in which open syntaxin provides from modest to robust rescue, open syntaxin uniquely aggravates the phenotypes of *unc-18* null mutants. Thus, KI open syntaxin exhibits a specific genetic interaction with loss-of-function mutants of *unc-18*.

### The open syntaxin-1 partially bypasses the requirement of Munc-13-1 for liposome fusion

Membrane fusion assays using reconstituted proteoliposomes provide a useful tool to correlate the physiological effects of mutations in components of the release machinery observed *in vivo* with the effects of these mutations *in vitro* on membrane fusion using minimal systems, yielding key insights into the mechanisms underlying the phenotypes ^4, 52^. Thus, the observation that fusion between liposomes containing synaptobrevin (V-liposomes) and liposomes containing syntaxin-1 and SNAP-25 (T-liposomes) in the presence of NSF and α-SNAP requires Munc18-1 and a Munc13-1 C-terminal fragment led to a model whereby Munc18-1 and Munc13-1 orchestrate SNARE complex assembly in an NSF-αSNAP-resistant manner, and provided a basis to understand the critical requirement of Munc18-1/UNC-18 and Munc13/UNC-13 for neurotransmitter release ^53^. Moreover, a P335A gain-of-function mutation in Munc18-1 (corresponding to P334A in *C. elegans unc-18*) partially bypassed the strict requirement of Munc13-1 for liposome fusion, mirroring the partial rescue of *unc-13* phenotypes by the P334A *unc-18* mutation observed *in vivo* ^45^.

We used an analogous approach to examine how the LE mutation that opens syntaxin-1 affects the requirement of Munc13-1 and Munc18-1 for liposome fusion, and thus test whether these reconstitution experiments correlate with the phenotypes of our genetic studies. Using an assay that simultaneously monitors lipid and content mixing ^54^, we observed fast, Ca^2+^-dependent fusion between V-liposomes and T-liposomes containing WT syntaxin-1 in the presence of NSF, α-SNAP, Munc18-1, a fragment spanning the C1, C2B, MUN and C2C domains of Munc13-1 (C1C2BMUNC2C) and a fragment spanning the C2 domains that form the cytoplasmic region of synaptotagmin-1 (C2AB) (Fig. 8a, 8b). Similar results were obtained in parallel experiments where the T-liposomes contained the open syntaxin-1 mutant, although we observed substantial Ca^2+^-independent fusion. Importantly, no fusion was observed in the absence of Munc18-1 regardless of whether the T-liposomes contained WT or open mutant syntaxin-1 (Fig. 8a, 8b). Moreover, considerable Ca^2+^-dependent fusion was observed in experiments performed with the syntaxin-1 open mutant where Munc18-1 was included but the Munc13-1 C1C2BMUNC2C fragment was absent, whereas practically no fusion was observed under these conditions with the liposomes containing WT syntaxin-1 (Fig. 8c, 8d, Fig. S7). These results are reminiscent of those obtained with the Munc18-1 P335A mutant ^45^ and another Munc18-1 gain-of-function mutant (D326K) ^55^, and correlate with the partial rescue of *unc-13* phenotypes by the syntaxin-1 open mutation. Overall, these data nicely correlate with our *in vivo* experiments showing that the open syntaxin mutant can partially rescue *unc-13* phenotypes but not the phenotypes of *unc-18* mutants, and are consistent with the notion that mutations that facilitate SNARE complex formation render membrane fusion and neurotransmitter release less critically dependent on Munc13/UNC-13.

**Figure 8.**
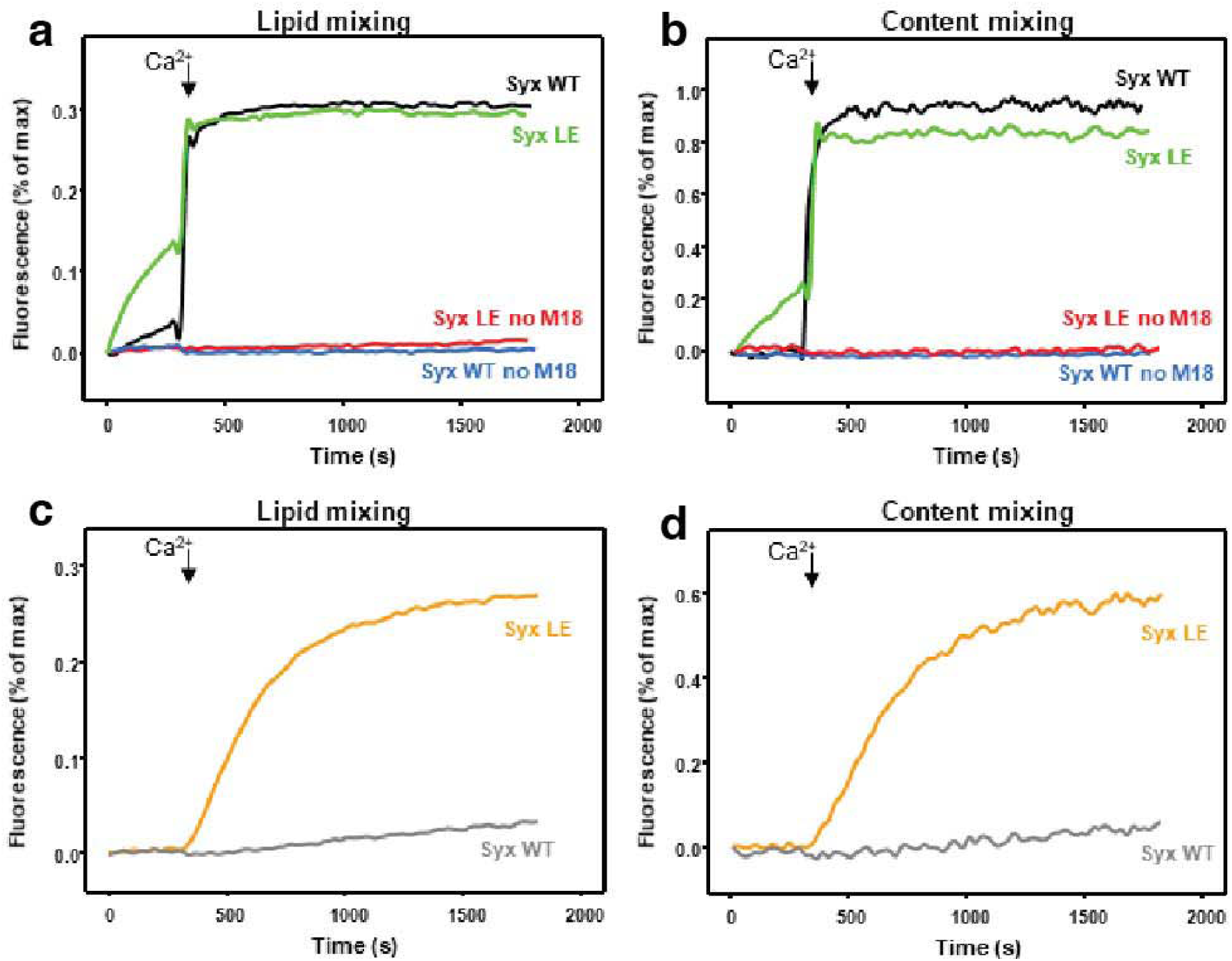
The syntaxin-1 open mutant partially bypasses the requirement of Munc13-1 but not Munc18-1 for liposome fusion. (a-d). Lipid mixing (a,c) between V- and T-liposomes (containing syntaxin-1 WT or LE mutant) was monitored from the fluorescence de-quenching of Marina Blue lipids and content mixing (b,d) was monitored from the increase in the fluorescence signal of Cy5-streptavidin trapped in the V-liposomes caused by FRET with PhycoE-biotin trapped in the T-liposomes upon liposome fusion. In A, B, assays were performed in the presence of NSF, αSNAP, the synaptotagmin-1 C2AB fragment and the Munc13-1 C1C2BMUNC2C fragment with (green and black traces) or without (red and blue traces) Munc18-1. In C, D, assays were performed in the presence of NSF, α SNAP, the synaptotagmin-1 C2AB fragment and Munc18-1, without the Munc13-1 C1C2BMUNC2C fragment. Experiments were started in the presence of 100 mM EGTA and 5 mM streptavidin, and Ca^2+^ (600 mM) was added at 300s.

### Open syntaxin animals do not significantly accelerate fusion pore kinetics of individual vesicles measured in the neuromuscular junction

Despite the availability of many synaptic mutants, most mutants do not affect the fusion kinetics of synaptic vesicle fusion. Importantly, a recent study of open syntaxin-1B in mice showed that it significantly reduces rise time in miniature EPSCs in high-fidelity Calyx of Held synapses ^32^, while this effect was not observed in more conventional hippocampal synapses ^31^. Therefore, we analyzed the kinetics of mPSCs of open syntaxin KI animals at the *C. elegans* neuromuscular junction. We found that there is no difference in rise time of mPSCs between wild-type control and open syntaxin KI animals (Fig. S8a-S8c). In addition to two excitatory acetylcholine receptors, one inhibitory GABA receptor functions at the *C. elegans* neuromuscular junction ^56^. The mPSC signals that we record are composed of both cholinergic mEPSCs and GABAergic mIPSCs and the mixture of two signals may interfere with the detection of subtle differences in fusion pore kinetics of individual vesicles.

To isolate cholinergic mEPSCs, we tested the effects of open syntaxin in the *unc-49(e407)* mutant, which lacks a functional GABA_A_ receptor. In the thrashing assay, *unc-49* null worms exhibited severely reduced motility (Fig. S9a). Interestingly, in the aldicarb assay, *unc-49* worms displayed hypersensitivity to aldicarb (Fig. S9b). We interpreted that the lack of GABA_A_ receptor contributes to the lack of relaxation of muscle activity which enhances the aldicarb-induced paralysis while interfering with the thrashing behavior. We found that the open syntaxin further increased aldicarb sensitivity of *unc-49* worms (Fig. S9b), again suggesting that open syntaxin can increase transmitter release in many different genetic backgrounds.

Electrophysiological recordings also indicated that open syntaxin induces a significant increase in frequency of cholinergic mEPSCs in the *unc-49(e407)* background (Fig. S9c, S9d). However, we did not find differences in the amplitude and rise times between *unc-49(e407)* mutants and *open syntaxin; unc-49(e407)* double mutants, as well as no differences in the kinetics of the mPSCs (Fig. S8d-S7f). Therefore, our results suggest that open syntaxin KI does not enhance fusion pore kinetics of synaptic vesicles to a detectable level in conventional synapses.

## Discussion

Great advances have been made in the past three decades in our understanding of the mechanism of neurotransmitter release, showing that release depends on SNARE complexes that bridge the vesicle and plasma membranes, and that SNARE complex assembly is orchestrated by Munc18-1/UNC-18 and Munc13/UNC-13 [reviewed in ^4^]. The tight complex between Munc18-1/UNC-18 and closed syntaxin serves as the starting point for this mechanism, and Munc13/UNC-13 was proposed to mediate the transition to the SNARE complex by opening syntaxin. A key finding that served as a basis for this model was that multicopy expression of the constitutively open LE syntaxin mutant ^18^ rescued the total abrogation of release observed in *C. elegans unc-13* nulls ^25^. Subsequently, KI of open syntaxin-1B in mice was shown to enhance spontaneous release and vesicular release probability, which was proposed to arise from enhanced SNARE complex assembly, although open syntaxin-1N KI did not rescue the lethal Munc13-1 KO phenotype ^31^.

Our data now show that KI of open syntaxin in *C. elegans* enhances exocytosis in an otherwise WT background as well as in a wide variety of genetic backgrounds characterized by diverse defects in synaptic transmission. These results indicate that the open syntaxin mutation provides a general means to enhance synaptic transmission and that increasing the number of SNARE complexes can overcome phenotypes that arise not only from defects in the SNARE complex assembly pathway but also from impairment of other aspects such of Ca^2+^ sensing, Ca^2+^ influx or neurotransmitter recognition. These findings also suggest that manipulating SNARE complex may provide a powerful tool for functional studies of synaptic transmission and neural networks, as well as opportunities to develop novel therapies for diverse neurological disorders.

The pioneering studies of Richmond et al. (2001) ^25^ provided crucial evidence demonstrating the critical functional importance of opening syntaxin in a physiological context. Although some of the conclusions might now be questioned, this work paved the way for a flourish of additional studies that investigated genetic interactions involving opening of syntaxin. The observation that multicopy expression of open syntaxin rescued not only *unc-13* but also of *unc-10* and *unc-31* phenotypes ^17, 30^ suggested that more than one protein might be involved in opening syntaxin and/or that Munc13/UNC-13 might have only an indirect role in this event. However, RIM/UNC-10 is known to play a key role in synaptic vesicle priming by activating Munc13-1/UNC-13 ^48^, and the functions of CAPS/UNC-31, which contains a MUN domain homologous to that of Munc13/UNC-13, are likely related to those of Munc13/UNC-13 [reviewed in ^57^]. Hence, these results were not incompatible with the proposed role of Munc13/UNC-13 in opening syntaxin.

Using an open syntaxin KI approach that does not suffer from technical problems that can arise from multicopy expression, our data now show that (i) *open syntaxin KI* increases synaptic vesicle exocytosis in an otherwise wild-type background (Fig. 1); (ii) *open syntaxin KI* can increase neurotransmitter release in a wide range of synaptic mutants that include *snt-1* (Figs. 2, 3), *unc-2* (Fig. 4, Fig. S3), *tom-1* (Fig. 6), *unc-31* (Fig. S4), and *unc-49* (Fig. S9), in addition to the previously reported *unc-13* (Fig. 5) and *unc-10* (Fig. S4); and (iii) the rescue of various *unc-13* mutants (Fig. 5) by open syntaxin KI is rather weak and contrasts with the stronger rescues observed for some of the other mutants. The gain-of-function caused by open syntaxin KI in *C*. *elegans* is reminiscent of the increased spontaneous release and vesicular release probability observed in open syntaxin-1B KI mice ^31^. Conversely, we did observe an increase in the charge transfer of evoked release in the open syntaxin KI worms (Fig. 3e) that was not observed in open syntaxin-1B KI mice ^31^. Further research will be needed to assess whether this difference arises from distinct impairment of the RRP, which was decreased in the KI mice ^31^. Regardless of this difference, the gain-of-function phenotypes caused by the open syntaxin mutation in mice and worms are most likely associated with the fact that this mutation facilitates SNARE complex assembly ^18^, and an increase in the number of assembled SNARE complexes enhances the probability of vesicle release ^32^. Although the importance of SNARE complex assembly for neurotransmitter release has been recognized for a long time, it is remarkable that open syntaxin can rescue the defects caused by ablation of proteins with functions as diverse as those of the Ca^2+^ sensor synaptotagmin, a Ca^2+^ channel and a GABA_A_ receptor.

The limited rescue of *unc-13* phenotypes by the open syntaxin KI (Fig. 5) correlates with studies that observed no rescue or much weaker rescues in *C. elegans unc-13* ^15, 58^ and in Munc13 deficient mice ^31, 34^ than those reported by Richmond et al. (2001). One might argue that such limited rescues arise because of the very strong nature of the *unc-13* phenotypes, which constitutes a strong challenge for any type of rescue. However, the rescue afforded by open syntaxin KI is rather modest even when combined with the *tom-1(ok285)* mutation (Fig. 5), which alleviates the severity of the *unc-13* phenotypes. Moreover, open syntaxin KI produces considerably stronger rescues of other mutants (e.g. *unc-2*, Fig. 4), and the P334A mutation in UNC-18 rescues the aldicarb sensitivity of the *unc-13(s69)* mutant better than open syntaxin (Fig. 6). Based on these findings, one might question whether Munc13/UNC-13 actually has a direct role in opening syntaxin. However, biochemical and biophysical data have shown that the Munc13-1 MUN domain accelerates the transition from the Munc18-1/closed syntaxin complex to the SNARE complex ^27^, most likely through direct interactions with syntaxin ^28, 29^. A plausible explanation for the apparent contradiction between these findings and the limited rescue of *unc-13* phenotypes by open syntaxin is that opening syntaxin is one of the roles of Munc13/UNC-13 but is not the primary function or at least not the sole primary function. In this context, the large, conserved C-terminal region of Munc13-1 has been shown to have a highly elongated structure ^59^ that has been proposed to provide a bridge between the plasma membrane and synaptic vesicles^54^. Strong evidence supporting this model has been recently provided by the observation that even a single point mutation in this 200 kDa protein that abolishes its membrane-membrane bridging ability abrogates neurotransmitter release almost completely ^60^. Note also that Munc13-1 has been shown to improve the fidelity of SNARE complex assembly, preventing antiparallel orientations of the SNAREs ^34^.

Interestingly, multicopy expression of open syntaxin cannot rescue *unc-18* phenotypes ^9^, and in our experiments with open syntaxin KI the phenotypes of all the mutant strains that we tested were improved except for *unc-18* mutants. In fact, open syntaxin KI aggravates *unc-18* phenotypes (Fig. 7, S5, S6). This lack of rescue in the absence of UNC-18 and the partial rescue in the absence of UNC-13 are nicely mirrored in our reconstitution assays, where the open syntaxin-1 mutant enables some liposome fusion in the absence of Munc13-1 C_1_C_2_BMUNC_2_C (unlike WT syntaxin-1), but not in the absence of Munc18-1 (Fig. 8). These findings can be readily explained by results from reconstitution assays showing that Munc18-1 and Munc13-1 are normally essential to form trans-SNARE complexes in the presence of NSF and αSNAP, because Munc18-1 and Munc13-1 orchestrate SNARE complex assembly in an NSF-αSNAP-resistant manner ^53, 54, 61^. The requirement of Munc13-1 for liposome fusion can be partially bypassed by the open syntaxin mutation (Fig. 8) or by mutations in Munc18-1 that also facilitate SNARE complex assembly ^45, 55^. However, Munc18-1 cannot be bypassed, likely for three reasons. First, because Munc18-1 provides a template to assemble the SNARE complex ^55, 62, 63^. Second, because αSNAP completely blocks SNARE-dependent liposome fusion through diverse interactions with the SNAREs and such inhibition can only be overcome by binding of Munc18-1 to closed syntaxin-1, suggesting that this complex constitutes an essential starting point for the productive pathway that leads to neurotransmitter release ^64^. Thus the observation that the open syntaxin mutation exacerbates rather than alleviates the *Unc18* phenotypes can be rationalized by the impairment of the interaction of syntaxin with Unc18/Munc18-1 caused by this mutation. And a third reason is that anterograde trafficking of syntaxin/UNC-64 is known to be severely impaired in *unc-18* mutants ^21^, and similar defects have been demonstrated in Munc18-1/2 double knockdown PC12 cells ^19, 20, 65, 66^ as well as Munc18-1 KO neurons ^22^. This trafficking defect is believed to be due to the formation of ectopic SNARE complexes in the ER and Golgi^67^. Such ectopic SNARE formation is accelerated when syntaxin/UNC-64 adopts the open conformation, which may contribute to the observed aggravation of *unc-18* phenotypes.

Does open syntaxin enhance fusion pore opening kinetics of individual vesicles? Several mechanisms to control fusion pore opening have been suggested, including the copy number of SNARE complexes. Nevertheless, there were almost no synaptic mutants known that affect the fusion kinetics of synaptic vesicle fusion. Using Calyx of Held synapses in mice, Acuna et al., demonstrated that open syntaxin-1B can reduce rise time of mEPSCs ^32^. The results appear to be consistent with a recent study showing that cleavage of synaptobrevin by tetanus toxin reduces the amplitude and kinetics of fusion events in cultured neurons ^68^. We did not observe changes in amplitude or kinetics of mEPSCs in open syntaxin KI despite a significant increase in frequency of mEPSCs. One potential explanation is that, in the Calyx of Held, the rise time is very fast (∼0.1 ms), which may imply that the diffusion time of neurotransmitter glutamates to reach receptors is short, allowing detection of small changes in the kinetics of vesicle fusion. The rise time in *C. elegans* neuromuscular junctions is significantly slower at ∼0.6 ms. However, the effects of synaptobrevin cleavage by tetanus toxin on the amplitude and kinetics were clearly observed in hippocampal cultured neurons in which the rise time is 1-2 ms ^68^. It might be possible to find synaptic mutants which can slow down the kinetics of vesicle fusion events at the appreciable level in *C. elegans* in the future.

Taken together, our results show that open syntaxin can suppress a wide range of exocytosis defects but not those of Munc18/UNC-18 mutants. Recent studies show that mutations in syntaxin-1B, Munc18, Munc13 and Tomosyn underlie a wide spectrum of childhood epilepsy and autism spectrum disorders (ASDs) ^69–77^. Similarly, mutations in proteins involved in the trafficking of postsynaptic neurotransmitter receptors such as Shank3 ^78–80^ and neuroligin-3 and -4 ^81^ are strongly implicated with ASD. Thus, small chemicals that facilitate opening of syntaxin-1 have the potential to alleviate many neurological diseases derived from defects in synaptic transmission by increasing synaptic transmitter release.

## Experimental Procedures

### CRISPR-mediated genome editing of unc-64

The strategy is shown in Fig. S1. First, a two-step PCR was performed to generate two different PCR products encoding two different guide RNA (sgRNAs) that would target g<u>CTGTACCTGCCTACAAGG</u>cgg sequence and <u>AAGACGAACCCAGAGAACAT</u>cgg sequence within the intron of *unc-64* genomic region (see Fig. S2). Second, a repair vector was constructed that contained a dual-marker selection cassette (GFP and neomycin-resistant gene ^35^ flanked by left and right homology arms of *unc-64*. Upstream (left arm) and downstream (right arm) sequences of the CRISPR target site were amplified by PCR using pTX21 (a kind gift from Dr. Michael Nonet), which contains the genomic sequence of the *unc-64* gene ^26^. The left arm PCR product (∼1.1 kb) was digested with SacI and NotI, while the right arm PCR product (∼1.7 kb) was digested with SpeI and ClaI. Each of the digested products was ligated into pBluescript, followed by sequencing of the ligated products. To generate a knock-in (KI) mutant allele, site-directed mutagenesis was conducted on pBluescript containing the right arm. The mutagenesis product was verified by sequencing. The dual-marker selection cassette (∼5.4 kb) digested with SpeI and NotI was ligated into pBluescript containing the both arms ^35^. Third, multiple plasmids were injected into N2 animals: a vector encoding Cas9, the repair vector, the DNA product of sgRNA, and several injection markers (P*myo-3-*mCherry and P*rab-3-*mCherry). Fourth, screening for knock-in animals began by applying neomycin to the progeny of the injected animals. Then, animals that were neomycin resistant, GFP positive, and mCherry negative were selected as knock-in candidates and genotyped for verification, resulting in the selection of *unc-64(sks3) worm*. We then injected Cre-recombinase gene into *unc-64(sks3)* worms to remove GFP and neomycin resistant gene. The genomic sequence of the resulting worm *unc-64(sks4)* was shown in Fig. S2.

### Genetics

All strains used in the study were maintained at 22°C on 30 mm agar nematode growth media (NGM) plates seeded with OP50, a strain of *E. coli*, as a food source. The *C. elegans* strains used were listed in Table 1. *unc-10(md1117)*, *unc-13(e51), (e1091), (n2813)*, *unc-31(e928)*, *unc-2(e55), unc-18(md299), snt-1(md290)* worms were purchased from the Caenorhabditis Genetics Center (University of Minnesota); VC223 strain with a genotype of *tom-1(ok285)* was provided by the *C. elegans* Reverse Genetics Core Facility at the University of British Columbia; *unc-2(hp647)* was generated in Mei Zhen’s lab ^42^. We crossed these mutants with *unc-64(sks4)* to generate double and triple mutants. Strains with *zxIs6* background were cultured in the dark at 22 °C on OP50-seeded NGM plates supplemented with all-trans retinal (0.5 mM). 15-20 adult worms from each plate were dissolved in worm lysis buffer to extract their DNA for PCR. PCR was conducted to confirm the genotype of double or triple mutants using primers purchased from IDT DNA.

**Table 1.** C. elegans knock-in mutants, alleles, and strains.

| Genotype | Strain # |
| --- | --- |
| *unc-64(sks3[L166A/E167A + loxP P<sub>myo-2</sub>-GFP; P<sub>rps-27</sub>-NeoR loxP) III* | UHN28 |
| *unc-64(sks4[L166A/E167A + loxP]) III* | UHN29 |
| *unc-2(e55) X; unc-64(sks4) III* | UHN30 |
| *unc-2(hp647) X; unc-64(sks4) III* | UHN31 |
| *unc-13(s69) I; unc-64(sks4) III* | UHN32 |
| *unc-13(e51) I; unc-64(sks4) III* | UHN33 |
| *unc-13(e0191) I; unc-64(sks4) III* | UHN34 |
| *unc-13(n2813) I; unc-64(sks4) III* | UHN35 |
| *tom-1(ok285) I; unc-64(sks4) III* | UHN36 |
| *unc-13(s69) tom-1(ok285) I; unc-64(sks4) III* | UHN37 |
| *unc-13(s69) tom-1(ok285) I; unc-18(sks2) X* | UHN26 |
| *unc-10(md1117) X; unc-64(sks4) III* | UHN38 |
| *unc-31(e928) IV; unc-64(sks4) III* | UHN39 |
| *snt-1(md290) II; unc-64(sks4) III* | UHN40 |
| *unc-18(md299) X; unc-64(sks4) III* | UHN41 |
| *unc-18(e81) X; unc-64(sks4) III* | UHN42 |
| *unc-18(sks1) X; unc-64(sks4) III* | UHN43 |
| *unc-49(e407) unc-64(sks4) III* | UHN44 |
| *unc-64(sks4) III; zxls6 V* | UHN45 |
| *snt-1(md290) II; zxls6 V* | UHN46 |
| *snt-1(md290) II; unc-64(sks4) III; zxls6 V* | UHN47 |

### Western blot analysis of UNC-64 expression in C. elegans worms

Protein extract was prepared as described previously by ^9^. Briefly, we cultured N2, *unc-64(sks3)* and *unc-64(sks4)* worms on 3.5 cm NGM plates When the plates were full of worms with little OP50 left, worms were harvested and washed three times with a buffer containing 360 mM sucrose and 12 mM HEPES. These worms were resuspended in ~ 5 times volume of the buffer and frozen at −80°C until use. The defrosted worms were sonicated on ice 10 times with a 5 s burst. The resulting lysate was centrifuged for 15 min to pellet the cuticle, nuclei, and other debris. After centrifugation, the supernatant (final protein concentrations were ~ 3–5 mg/ml) was transferred to a clean microcentrifuge tube with an equal volume of sample buffer (2X). Fifty micrograms of samples were subjected to SDS-PAGE electrophoresis followed by immunoblotting. The expression of UNC-64 was detected by polyclonal anti-syntaxin-1 antibody (I378, 1:1000) ^18^.

### Behavioral analyses

Motility of each strain was determined by counting the thrashing rate of *C. elegans* in liquid medium. Briefly, worms were bleached to release their eggs and were then synchronously grown to young adulthood. Young adult worms were placed in a 60 μL drop of M9 buffer on a 30 mm petri dish cover. After a 2-minute recovery period, worms were video-recorded for 2 minutes using an OMAX A3580U camera on a dissecting microscope with the OMAX ToupView software. The number of thrashes per minute were manually counted and averaged within each strain. A thrash was defined as a complete bend in the opposite direction at the midpoint of the body. At least 40 worms for each strain was measured in each analysis.

### Aldicarb assays

Aldicarb sensitivity was assessed using synchronously grown adult worms placed on non-seeded 30mm NGM plates containing 0.3 mM or 1 mM aldicarb. All assays were done in 1 mM aldicarb plates unless specified otherwise. Over a 4 or 24-hour period, worms were monitored for paralysis at 15- or 30-min intervals. Worms were considered paralyzed when there was no movement or pharyngeal pumping in response to 3 taps to the head and tail with a platinum wire. Once paralyzed, worms were removed from the plate. 6 sets of 10-20 worms were examined for each strain.

### Growth speed analyses

Growth speed was assessed by eye using an OMAX A3580U camera on a dissecting microscope. Briefly, 15 eggs were extracted from adult worms into OP50-seeded NGM plates and placed along the edge of the bacterial lawn roughly equidistance from each other. Photos of the plates were taken daily at 1X magnification with the OMAX ToupView software for up to 7 days for N2 worms, and up to 9 days for the other strains. Growth speed and body size were compared by eye based on photos taken on day 9.

### Electrophysiology

The dissection of the *C. elegans* was described previously (Gao and Zhen, 2011, Richmond et al., 1999). Briefly, one- or two-days old hermaphrodite adults were glued to a sylgard (Dow Corning, USA)-coated cover glass covered with bath solution. The integrity of the neuromuscular junction preparation was visually examined via DIC microscopy, and anterior muscle cells were patched using fire-polished 4−6 MΩ resistant borosilicate pipettes (World Precision Instruments, USA). Membrane currents were recorded in the whole-cell configuration by HEKA EPC-9 patch clamp amplifier, using the PULSE software and processed with Igor Pro 6.21 (WaveMetrics) and Clampfit 10 (Axon Instruments, Molecular Devices, USA). Data were digitized at 10 kHz and filtered at 2.6 kHz.

Light stimulation of *zxIs6* was performed with an LED lamp at a wavelength of 460 ± 5 nm (3.75 mW/mm^2^), triggered by the PULSE software for 10 ms. Muscle cells were recorded with a holding potential of 60 mV. The recording solutions were as described in our previous L studies ^82^. Specifically, the pipette solution contains (in mM): K-gluconate 115; KCl 25; CaCl_2_ 0.1; MgCl_2_ 5; BAPTA 1; HEPES 10; Na_2_ATP 5; Na_2_GTP 0.5; cAMP 0.5; cGMP 0.5, pH 7.2 with KOH, ∼320 mOsm. The bath solution consists of (in mM): NaCl 150; KCl 5; CaCl_2_ 5; MgCl_2_ 1; glucose 10; sucrose 5; HEPES 15, pH 7.3 with NaOH, ∼330 mOsm. All chemicals were from Sigma. Experiments were performed at room temperatures (20−22℃).

### Protein expression and purification

Bacterial expression and purification of full-length rat syntaxin-1A, the rat syntaxin L165E/E166A (LE) mutant, a cysteine-free variant of full-length rat SNAP-25a, a full-length rat synaptobrevin-2, full-length rat Munc18-1, rat synaptotagmin-1 C2AB fragment (residues 131-421), full-length Chinese hamster NSF, full-length Bos Taurus α C1C2BMUNC2C fragment (residues 529-1725, Δ 408-1452) were described previously [and references therein]. We note that full-length rat syntaxin-1A was purified in buffer containing dodecylphosphocoline to prevent its aggregation ^84^.

### Lipid and content mixing assays

Assays that simultaneously measure lipid and content mixing were performed as previously described ^54^. Briefly, V-liposomes with reconstituted rat synaptobrevin-2 (protein-to-lipid ratio, 1:500) contained 39% POPC, 19% DOPS, 19% POPE, 20% cholesterol, 1.5% NBD PE, and 1.5% Marina Blue DHPE. T-liposomes with reconstituted rat syntaxin-1A or syntaxin-1 LE mutant together with rat SNAP-25A (protein-to-lipid ratio, 1:800) contained 38% POPC, 18% DOPS, 20% POPE, 20% cholesterol, 2% PIP2, and 2% DAG. Lipid solutions were then mixed with respective proteins and with 4 μ M Cy5-

Streptavidin for V-liposomes in 25 mM HEPES, pH 7.4, 150 mM KCl, 1mM TCEP, 10% glycerol (v/v). V-liposomes (0.125 mM lipids) were mixed with T-liposomes (0.25 mM lipids) in a total volume of 200 μL in the presence of 2.5 mM MgCl2, 2 mM ATP, 0.1 mM EGTA, 5μM streptavidin, 0.4 μM NSF, 2 μM α-SNAP, 1 μM Mync13-1, 1 μM synaptotagmin-1 C2AB and 1μμ M Munc13-1 C1C2BMUNC2C. Before mixing, T liposomes were incubated with NSF, MgCl_2_, ATP, EGTA, streptavidin, NSF, α Munc18-1 at 37°C for 25 min. 0.6 mM of CaCl_2_ was added at 300s to each reaction mixture. A PTI spectrofluorometer was used to measure lipid mixing from de-quenching of the fluorescence of Marina Blue–labeled lipids (excitation at 370 nm, emission at 465 nm) and content mixing from the development of FRET between PhycoE-Biotin trapped in the T-liposomes and Cy5-streptavidin trapped in the V-liposomes (PhycoE-biotin excitation at 565 nm, Cy5-streptavidin emission at 670 nm). All experiments were performed at 30°C. Lipid and content mixing were normalized as the percentage values of the maximum signals obtained by addition of 1% b-OG at the end to each reaction mixture (for lipid mixing) or to controls without streptavidin to measure maximal Cy5 fluorescence (for content mixing).

### Statistical analyses

Statistical analyses were done in OriginPro2018 using the independent t-test for two-group experiments with a *p* value less than 0.05 as the threshold for statistical significance. For comparison of multiple groups, one-way ANOVA (analysis of variance) was conducted, followed by Tukey’s range test, with a significance level of 0.05.

## Author Contributions

S.S., S.G., and J.R. designed and supervised experiments, analyzed data and wrote the manuscript. C-W.T., B.Y., and K.P.S performed experiments, analyzed data, prepared figures and wrote the manuscript. M.H. performed experiments and analyzed data. K.S. and X.X. contributed to the experiments. M.Z. provided the critical expertise and advice for the experiments and helped with editing the manuscript. P.P.M. and L.H. helped with editing the manuscript.

## Acknowledgements

This research was supported by the *Natural Sciences and Engineering Research Council* of Canada (RGPIN-2015-06438, to S.S.), the Canadian Institute of Health Research (MOP-130573, to S.S.), the Welch Foundation (I-1304 to J.R.) and the NIH (R35 NS097333 to J.R.), the National Natural Science Foundation of China (31871069, 31671052 to S.G.), the Junior Thousand Talents Program of China, and funds from Huazhong University of Science and Technology (Dengfeng Initiative, Global Talents Recruitment Program). *unc-10(md1117)*, *unc-13(e51), (e1091), (n2813)*, *unc-31(e928)*, *unc-2(e55), unc-18(md299) (e81), snt-1(md290)* worms were provided from the Caenorhabditis Genetics Center (University of Minnesota). VC223 strain with a genotype of *tom-1(ok285)* was provided by the *C. elegans* Reverse Genetics Core Facility at the University of British Columbia, which is part of the international *C. elegans* Gene Knockout Consortium. pTX21 plasmid and anti-syntaxin-1 polyclonal antibody are kind gifts from Dr. Michael Nonet (Washington University) and Dr. Thomas C. Südhof (Stanford University), respectively. We also thank Dr. Thomas Südhof for comments on an earlier version of the draft. The summer scholarship from Undergraduate Research Opportunity Program (UROP) from University of Toronto was awarded to C-W.T. in 2016 and 2017.

---

**Supplementary Figure S1.**
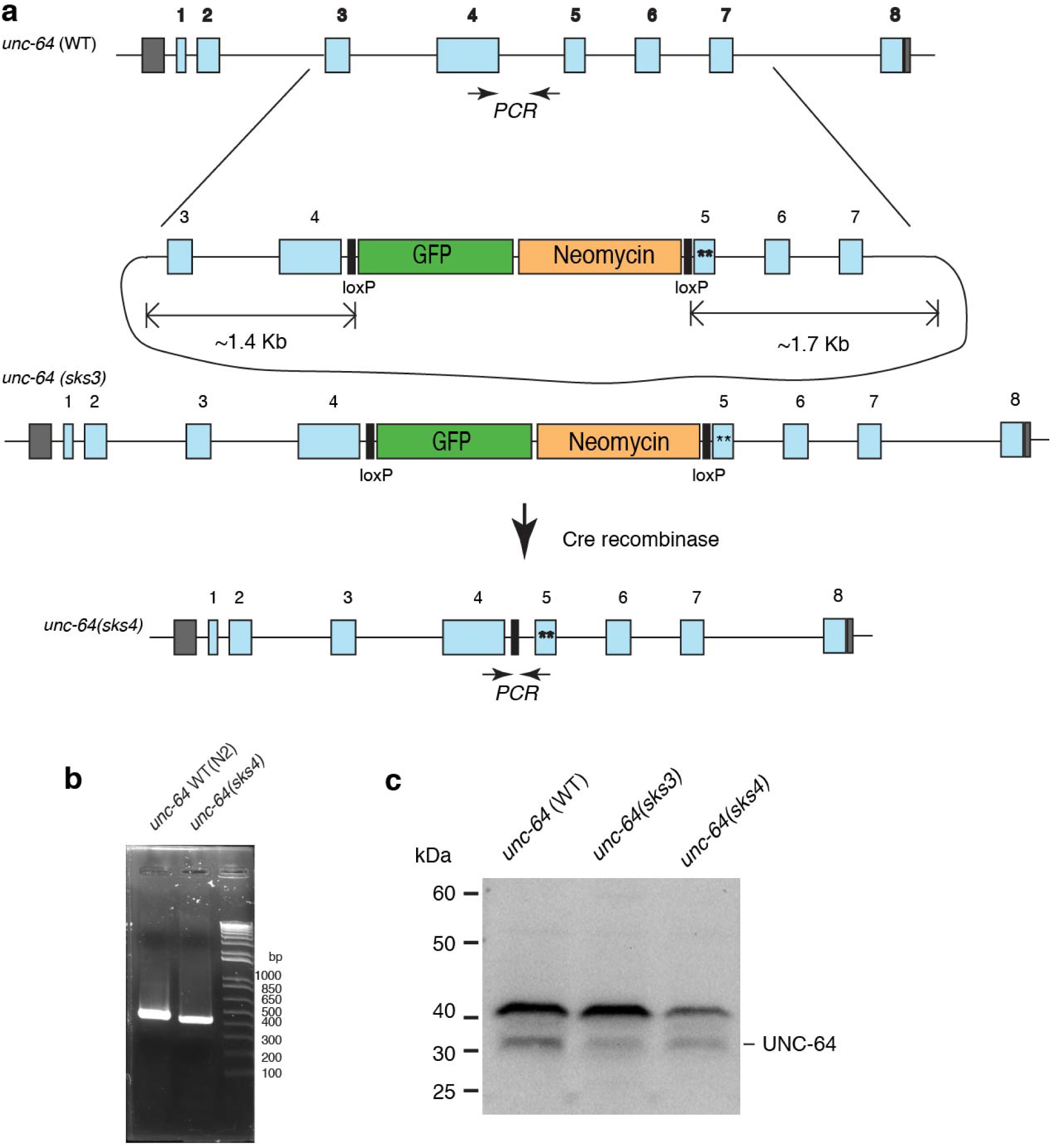
Generation of open syntaxin KI animals. (a) A schematic diagram represents how CRISPR-mediated homologous recombination replaces the wild-type allele of *unc-64* with the L166A/E167A mutant allele. (b) PCR (primers: GGTGTAAGGGACGAATTCAGAG, CAAACCTGTTGGCTATCTGTGA)-based genotyping of *unc-64(sks4)* LE open mutant allele in comparison with WT. (c) Fifty micrograms of indicated samples were subjected to SDS-PAGE followed by immunoblotting. The expression of UNC-64 protein was detected anti-syntaxin-1 polyclonal antibody (I378) ^18^ followed by enhanced chemiluminescence detection.

**Supplementary Figure S2.**
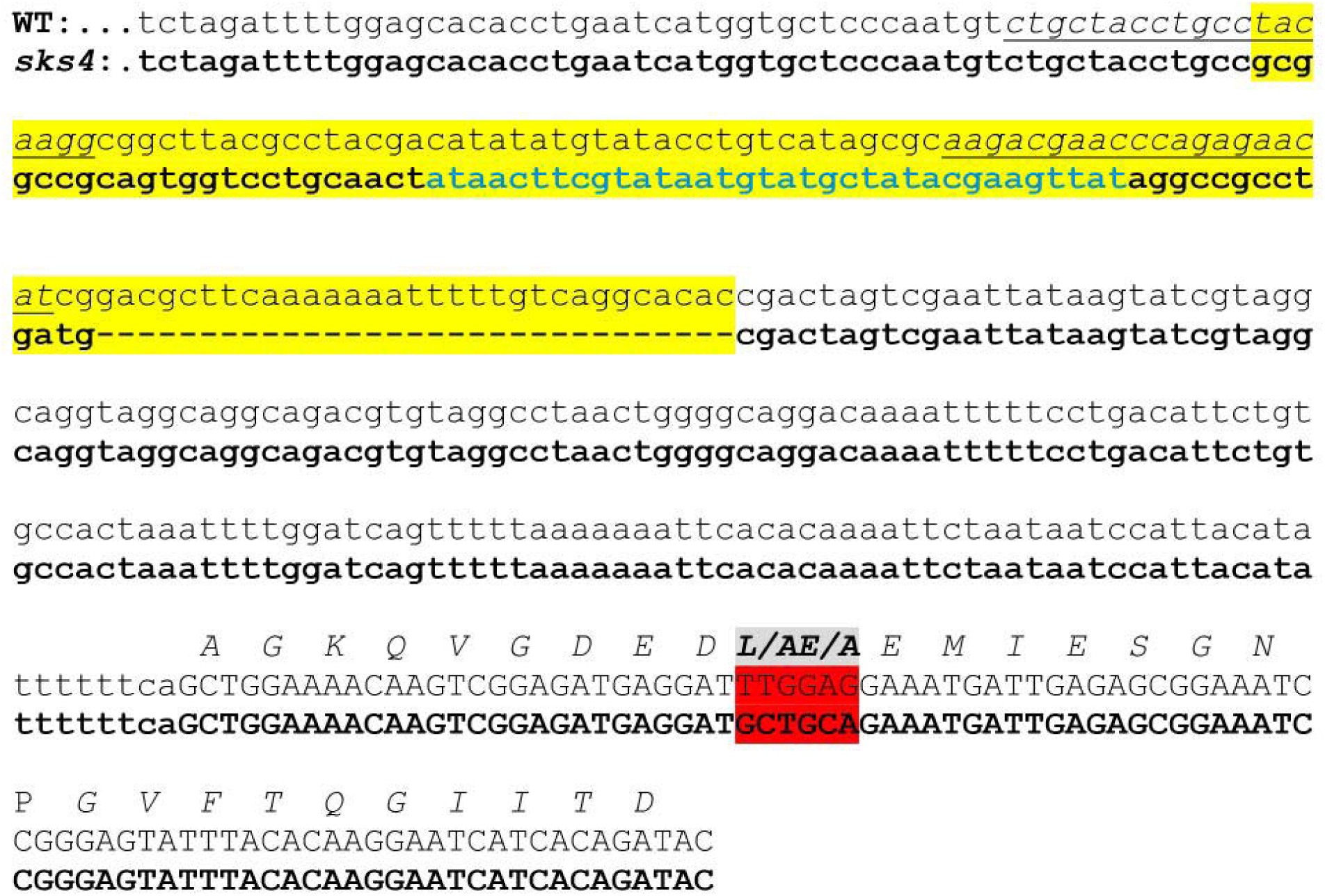
Comparison of genomic *unc-64* gene (WT allele from N2 vs. sks4 allele) surrounding the L166A/E167A mutations (highlighted by red). Lower letters indicate intron while capital letters indicate exon of *unc-64* genome. Different sequences between the two in the intron region were highlighted by yellow. Blue letters indicate loxP sequence. Underlines indicated the target sequences of Cas9.

**Supplementary Figure S3.**
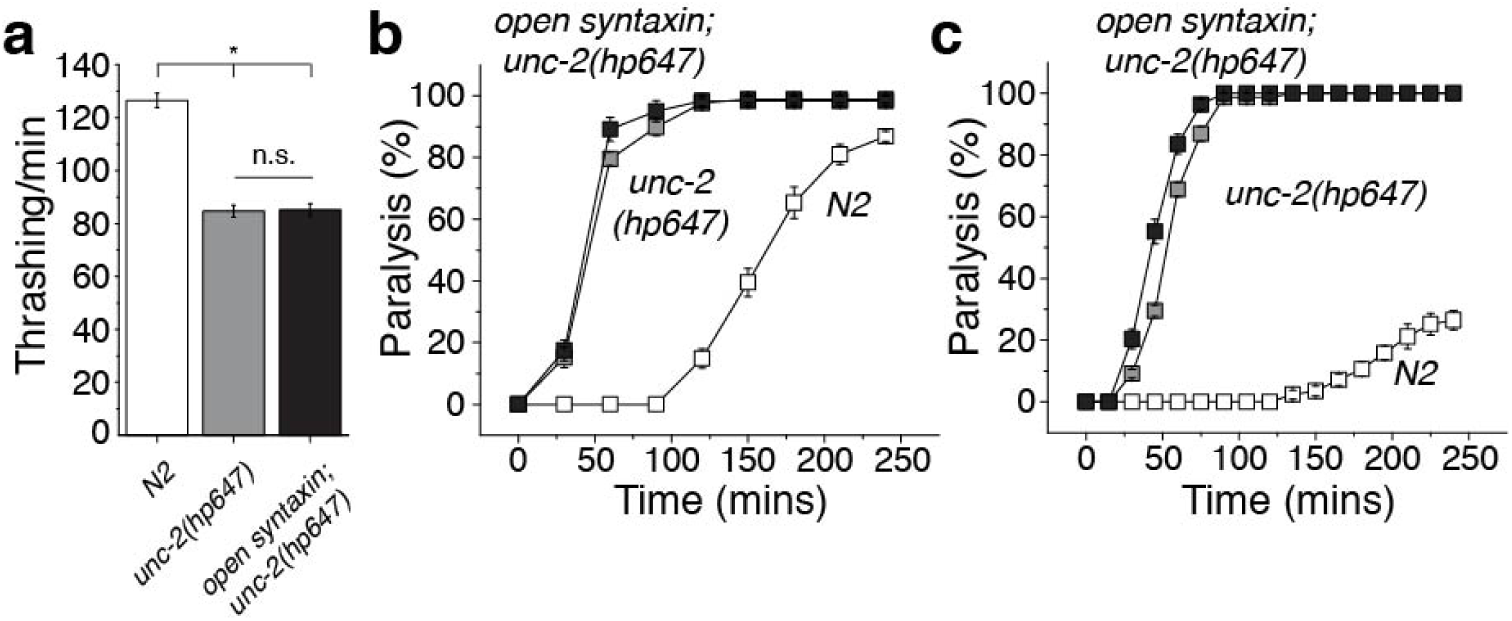
Open syntaxin enhances exocytosis in gain-of-function calcium channel mutants. (a) Thrashing assay of N2, *unc-2(hp647),* and *open syntaxin; unc-2(hp647)* double mutants in M9 buffer. Interestingly, *unc-2(hp647)* worms displayed reduced thrashing rates (84.7 thrashes/min) compared to N2 wild-types, which was unaffected by the introduction of open syntaxin KI in the double mutant (85.2 thrashes/min). n=40 for each strain. In one-way ANOVA statistical tests, *F*_(2,117)_ = 92.5 and *p =* 0. Tukey’s test was performed for means analysis in ANOVA. Error bars represent SEM. \**p* < 0.05. (b) Aldicarb assays of N2, *unc-2(hp647),* and *open syntaxin; unc-2(hp647)* double mutants. *unc-2(hp647)* and *open syntaxin; unc-2(hp647)* animals displayed similar hypersensitivity to 1mM aldicarb. n=6. Each assay was conducted with 15-20 worms. Error bars represent SEM. (c) Aldicarb assays of N2, *unc-2(hp647),* and *open syntaxin; unc-2(hp647)* double mutants on 0.3 mM aldicarb plates and probed for paralysis every 15 mins. The *open syntaxin; unc-2(hp647)* animals displayed slightly greater hypersensitivity to aldicarb. n=6. Each assay was conducted with 15-20 worms. Error bars represent SEM.

**Supplementary Figure S4.**
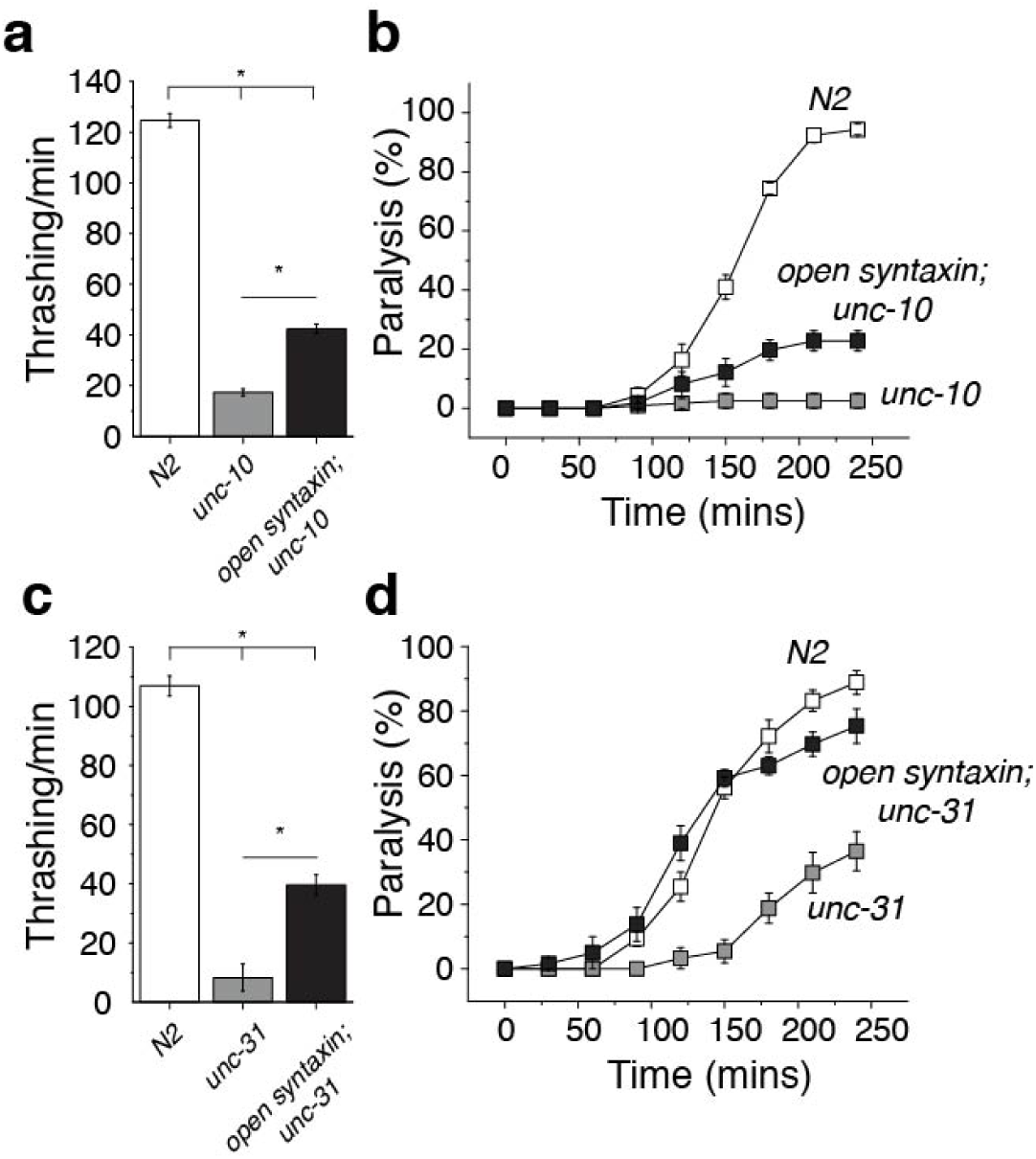
Open syntaxin partially rescues exocytosis and motility of *unc-10* and *unc-31* mutants. (a) Thrashing assay of N2, *unc-10(md1117),* and *open syntaxin; unc-10(md1117)* double mutants in M9 buffer. *unc-10(md1117)* worms displayed greatly reduced thrashing rates (17.3 thrashes/min) which was moderately increased by the introduction of open syntaxin in the double mutant (42.5 thrashes/min). n=40 for each strain. In one-way ANOVA statistical tests, *F*_(2,117)_ = 726 and *p =* 0. Tukey’s test was performed for means analysis in ANOVA. Error bars represent SEM. \**p* < 0.05. (b) Aldicarb assays of N2, *unc-10,* and *open syntaxin; unc-10(md1117)* double mutants. *open syntaxin; unc-10(md1117)* animals displayed slightly increased sensitivity to aldicarb compared to *unc-10(md1117)* single mutants. n=6. Each assay was conducted with 15-20 worms. Error bars represent SEM. (c) Thrashing assay of N2, *unc-31(e928),* and *open syntaxin; unc-31(e928)* double mutants in M9 buffer. *unc-31(e928)* worms displayed impaired thrashing rates (8.31 thrashes/min) which was increased by the introduction of open syntaxin in the double mutant (39.6 thrashes/min). n=40 for each strain. In one-way ANOVA statistical tests, *F*_(2,117)_ = 281 and *p =* 0. Tukey’s test was performed for means analysis in ANOVA. Error bars represent SEM. \**p* < 0.05. (d) Aldicarb assays of N2, *unc-31(e928),* and *open syntaxin; unc-31(e928)* double mutants. *open syntaxin; unc-31(e928)* animals displayed increased sensitivity to aldicarb to near wild-type levels compared to *unc-31(e928)* single mutants. n=6. Each assay was conducted with 15-20 worms. Error bars represent SEM.

**Supplementary Figure S5.**
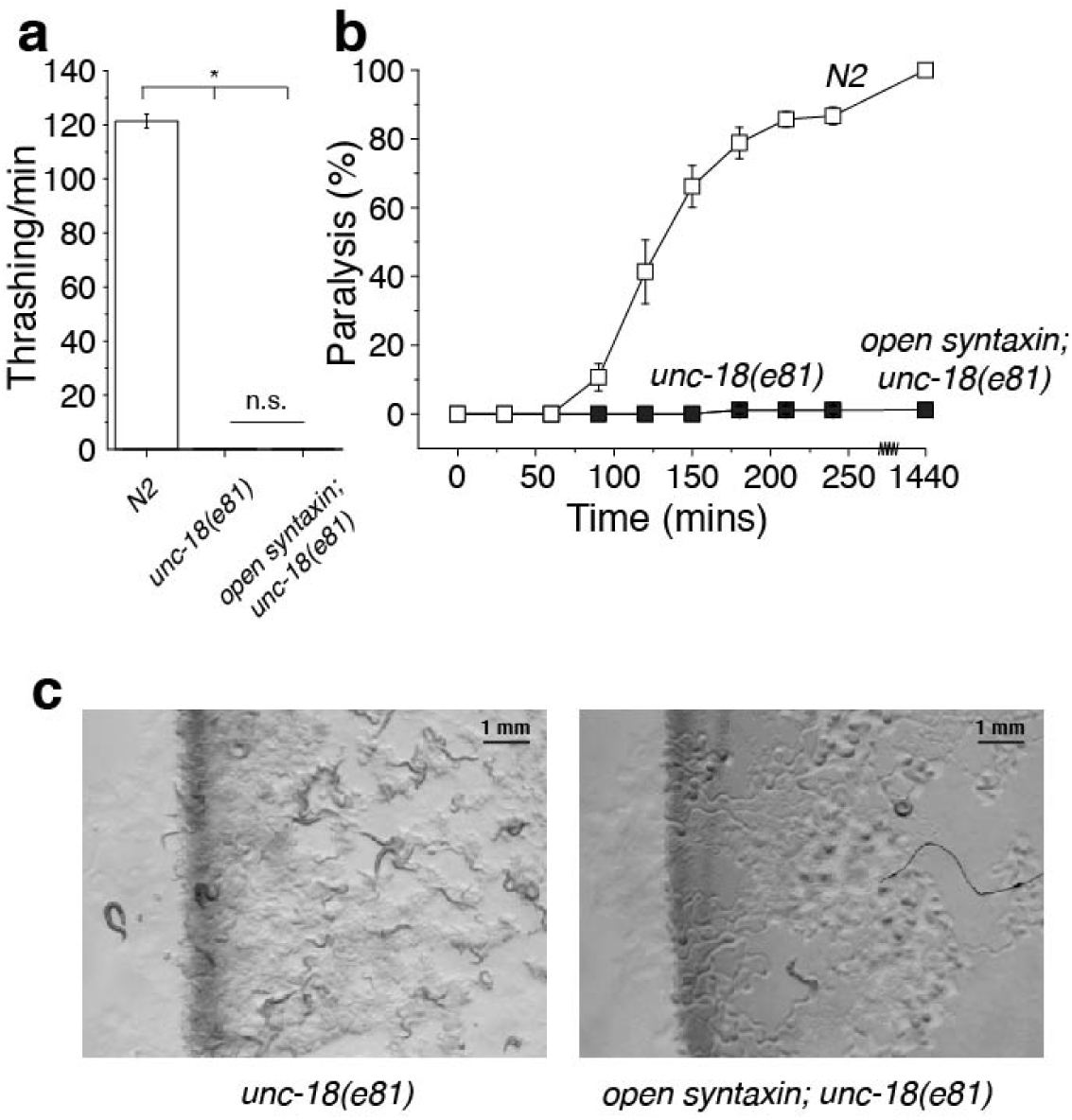
The knock-in open syntaxin mutation further impairs phenotypes of *unc-18(e81)* mutants. (a) Thrashing assay of N2, *unc-18(e81),* and *open syntaxin; unc-18(e81)* double mutants in M9 buffer. *unc-18(e81)* worms displayed greatly reduced thrashing rates (0.200 thrashes/min) which was reduced further by the introduction of open syntaxin in the double mutant to 0.100 thrashes/min. n=40 for each strain. In one-way ANOVA statistical tests, *F*_(2,117)_ = 2170 and *p =* 0. Tukey’s test was performed for means analysis in ANOVA. Error bars represent SEM. \**p* < 0.05; n.s. = not significant. (b) Aldicarb assays of N2, *unc-18(e81),* and *open syntaxin; unc-18(e81)* double mutants. *unc-18* and *open syntaxin; unc-18* animals displayed similar resistance to aldicarb even after 24 hours. n=6 for 4-hr assays, n=1 for 24-hr assay. Each assay was conducted with 15-20 worms. Error bars represent SEM. (c) Growth rate/body size images taken at day 9.

**Supplementary Figure S6.**
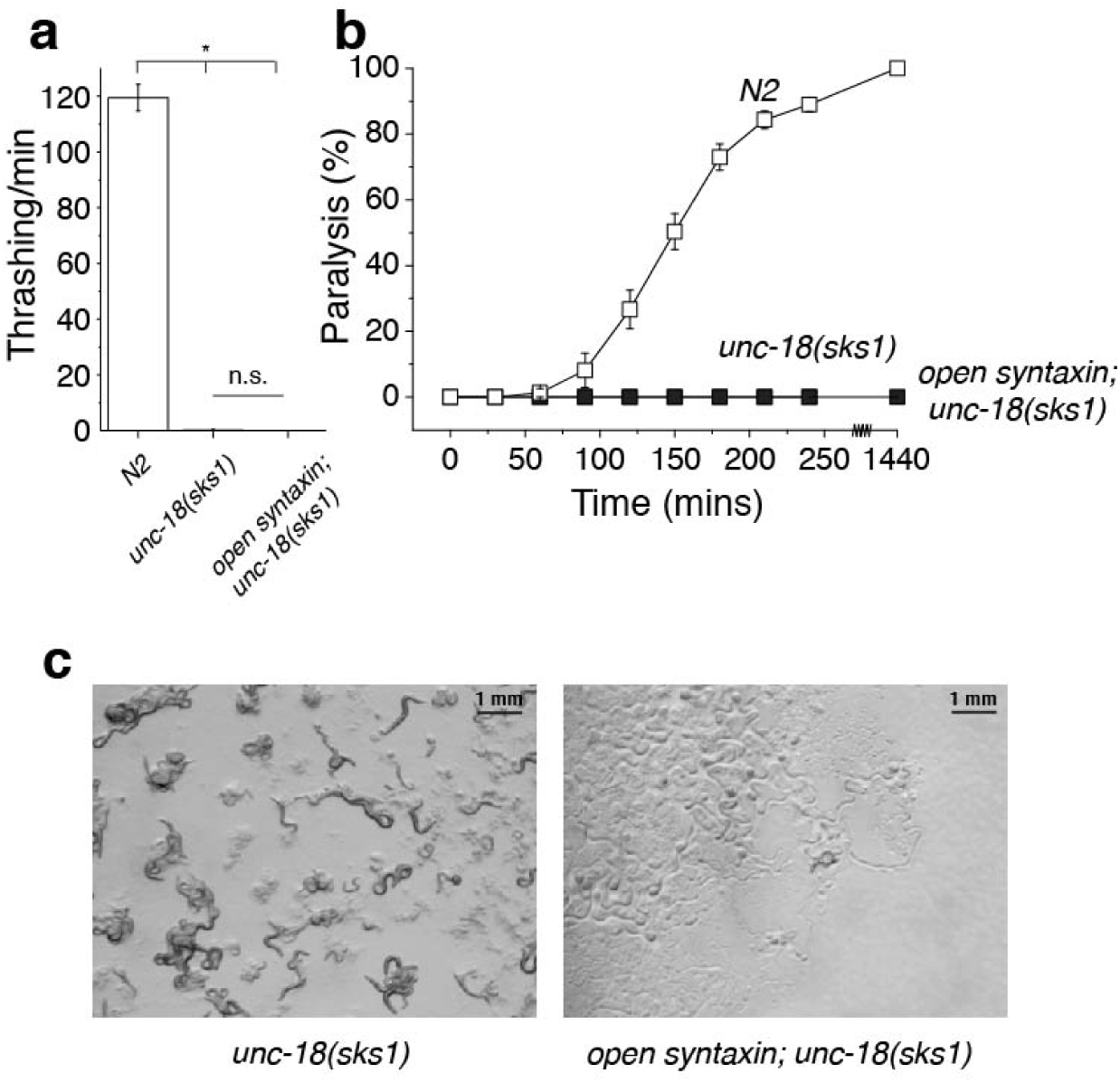
The knock-in open syntaxin mutation further impairs phenotypes of *unc-18(sks1)* mutants. (a) Thrashing assay of N2, *unc-18(sks1),* and *open syntaxin; unc-18(sks1)* double mutants in M9 buffer. *unc-18(sks1)* worms displayed greatly reduced thrashing rates (0.175 thrashes/min) which was reduced further by the introduction of open syntaxin in the double mutant to 0.0375 thrashes/min. n=40 for each strain. In one-way ANOVA statistical tests, *F*_(2,117)_ = 607 and *p =* 0. Tukey’s test was performed for means analysis in ANOVA. Error bars represent SEM. \**p* < 0.05; n.s. = not significant. (b) Aldicarb assays of N2, *unc-18(sks1),* and *open syntaxin; unc-18(sks1)* double mutants. *unc-18* and *open syntaxin; unc-18* animals displayed similar resistance to aldicarb even after 24 hours. n=6 for 4-hr assays. Each assay was conducted with 15-20 worms. Error bars represent SEM. (c) Growth rate/body size images taken at day 9.

**Supplementary Figure S7.**
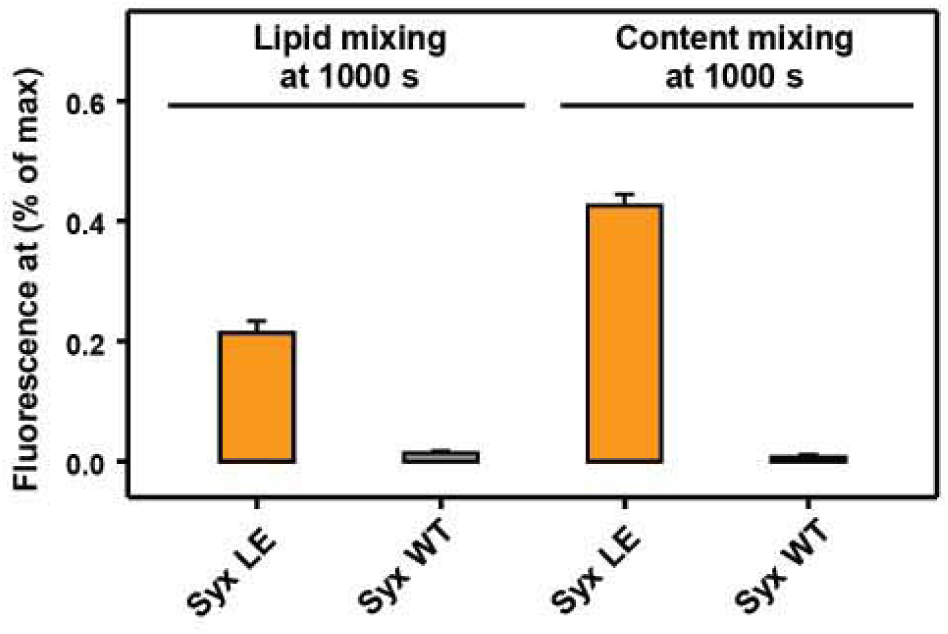
Quantification of the fusion assays shown in Fig. 8c, 8d. Bars represent averages of the normalized fluorescence intensities observed in lipid mixing (a) and content mixing (b) assays at 1,000 s, performed in triplicates. Error bars represent standard deviations.

**Supplementary Figure S8.**
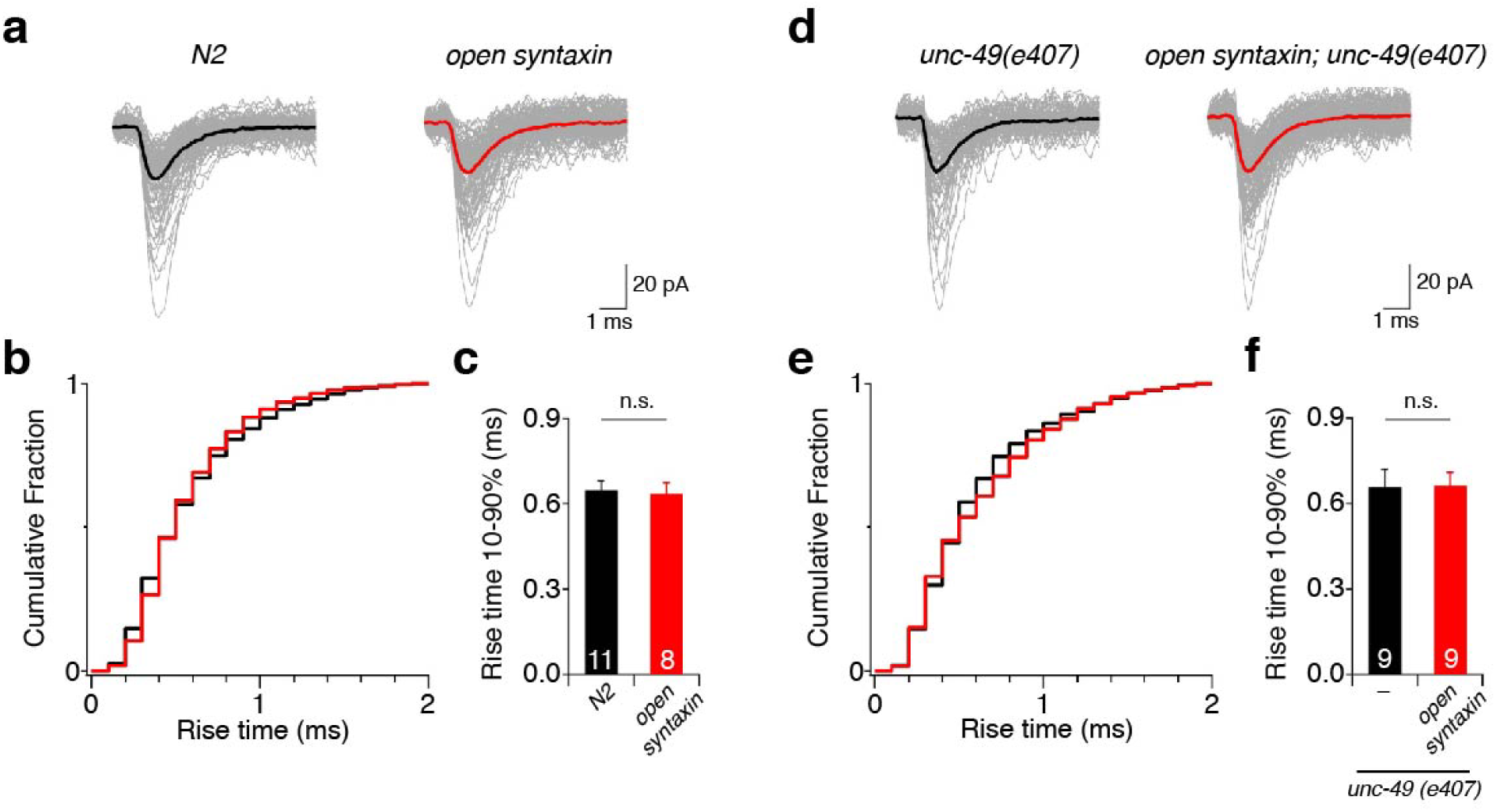
The open syntaxin knock-in mutation does not affect synaptic vesicle fusion pore kinetics appreciably. (a) Representative traces of mPSCs recorded from N2 and open syntaxin KI worms. (b). Cumulative distributions of mPSC rise times in N2 wild-type (black) and open syntaxin KI (red) worms. (c) Quantitative representation of 10-90% rise times in B. n.s. = not significant. n=11 for N2 and n=8 for *open syntaxin* animals. (d). Representative traces of mEPSCs recorded from *unc-49(e407)* and *open syntaxin; unc-49(e407)* double mutant worms. (e). Cumulative distributions of mEPSC rise times in *unc-49* (black) and *open syntaxin; unc-49* (red) worms. (f) Quantitative representation of 10-90% rise times in E. n.s. = not significant. n=9 for *unc-49(e407)* and *open syntaxin; unc-49(e407)* animals.

**Supplementary Figure S9.**
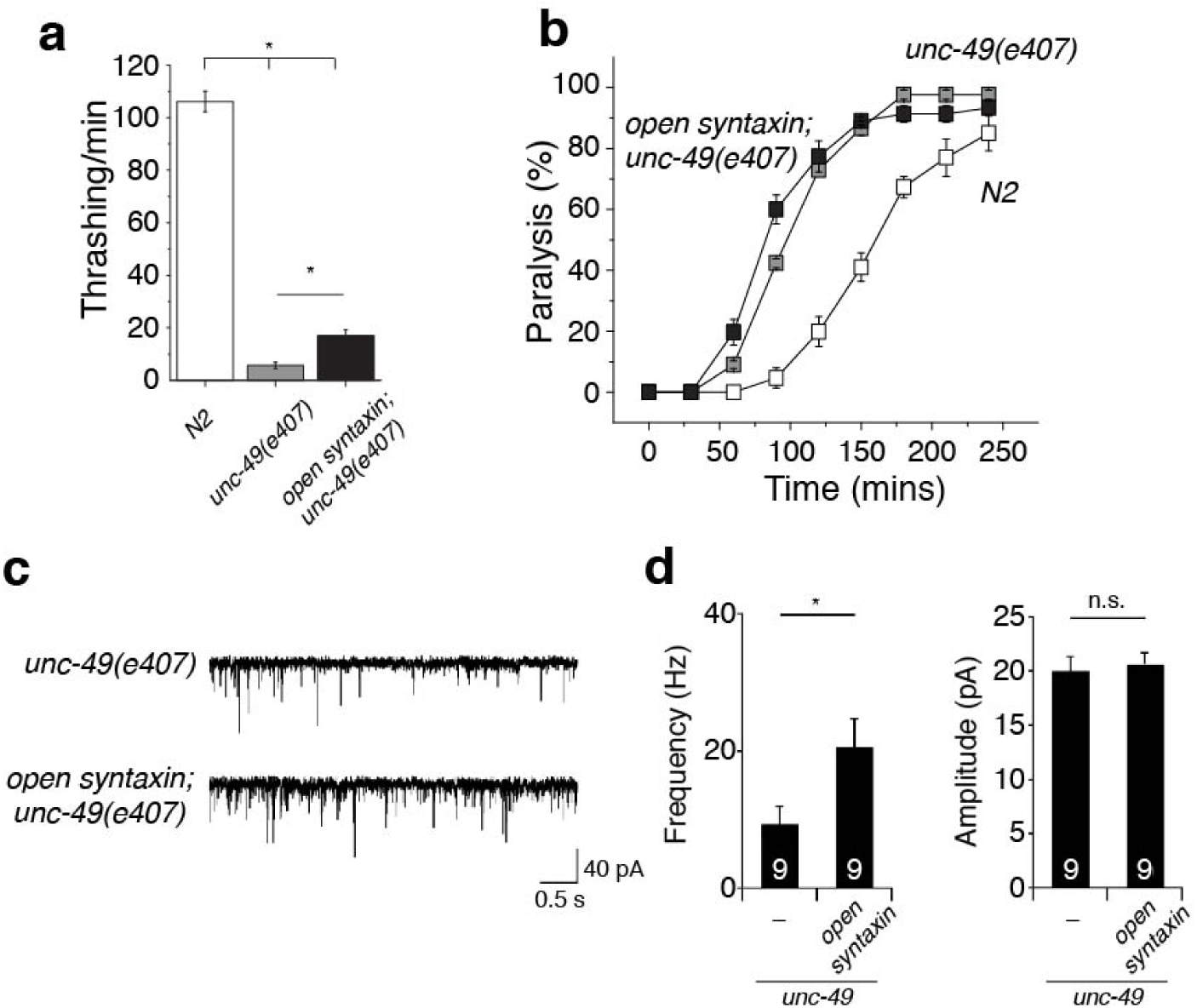
The knock-in open syntaxin mutation enhances exocytosis in GABA receptor mutants. (a) Thrashing assay of N2, *unc-49(e407),* and *open syntaxin; unc-49(e407)* double mutants in M9 buffer. *unc-49(e407)* worms displayed greatly reduced thrashing rates (5.73 thrashes/min) which was rescued by the introduction of open syntaxin in the double mutant (44.5 thrashes/min). n=40 for each strain. In one-way ANOVA statistical tests, *F*_(2,117)_ = 326 and *p =* 0. Tukey’s test was performed for means analysis in ANOVA. Error bars represent SEM. \**p* < 0.05. (b) Aldicarb assays of N2, *unc-49(e407),* and *open syntaxin; unc-49(e407)* double mutants. *unc-49* animals displayed hypersensitivity to aldicarb (grey), which was further enhanced in the double mutant (black). n=6. Each assay was conducted with 15-20 worms. Error bars represent SEM. (c) Representative mEPSC traces recorded from N2, *unc-49(e407),* and *open syntaxin; unc-49(e407)* worms. (d) Summary data of mEPSC frequency (left) and amplitude (right) of N2, *unc-49(e407),* and *open syntaxin; unc-49(e407)* worms. \**p* < 0.05; n.s. = not significant. n=9 for *unc-49(e407)* and *open syntaxin; unc-49(e407)* animals.

